# Understanding water conservation vs. profligation traits in vegetable legumes through a physio-transcriptomic-functional approach

**DOI:** 10.1101/2022.10.01.510433

**Authors:** Pingping Fang, Ting Sun, Arun Kumar Pandey, Libo Jiang, Xinyang Wu, Yannan Hu, Shiping Cheng, Xiaofang Li, Pei Xu

## Abstract

Vegetable soybean and cowpea are related warm-season legumes showing contrasting leaf water use behaviors under similar root drought stresses, whose mechanisms are not well understood. Here we conducted an integrative phenomic-transcriptomic study on the two crops grown in a feedback irrigation system that enabled precise control of soil water contents. Continuous transpiration rate monitoring demonstrated that cowpea used water more conservatively under earlier soil drought stages, but tended to maintain higher transpiration under prolonged drought. Interestingly, we observed a soybean-specific transpiration rate increase accompanied by phase shift under moderate soil drought. Time-series transcriptomic analysis suggested a dehydration avoidance mechanism of cowpea at early soil drought stage, in which the *VuHAI3* and *VuTIP2;3* genes were suggested to be involved. Multifactorial gene clustering analysis revealed different responsiveness of genes to drought, time of day and their interactions between the two crops, which involved species-dependent regulation of the core clock genes. Gene network analysis identified two co-expression modules each associated with transpiration rate in cowpea and soybean, including a pair of negatively-correlated modules between species. Module hub genes, including the ABA-degrading gene *GmCYP707A4* and the trehalose-phosphatase/synthase gene *VuTPS9* were identified. Inter-modular network analysis revealed putative co-players of the hub genes. Transgenic analyses verified the role of *VuTPS9* in regulating transpiration rate under osmotic stresses. These findings propose that species-specific transcriptomic reprograming in leaves of the two crops suffering similar soil drought was not only a result of the different drought resistance level, but a cause of it.

## Introduction

By growing under environmental conditions with a wide range of variations in water availability, plants develop flexible water use strategies in response to water scarcity. As the gateway of both water efflux and CO_2_ intake, stomatal control is critical for balancing drought tolerance and growth under soil water shortages^1^. Some plants develop a sensitive response to drought at the stomata level, representing a conservative water use strategy; conversely, some other plants adopt a “profligate” water use strategy that is marked by less sensitive stomatal control that allows transpiration of more water^2^. From an agronomic point of view, both strategies can be adaptive depending on the specific soil drought scenarios. Numerous studies have analyzed water use traits in crops including legumes, grains and trees^3-5^, yet current mechanistic understanding of the trait plasticity is still limited.

Cross-species comparison of the dynamic water use traits is often challenged by the difficulty of generating homogenous progressive soil stress in a reasonably long term^6^. Recently, several high-throughput platforms have emerged, some being equipped with feedback irrigation systems, to enable precise control of soil water content in each unit^7, 8^. One of such platforms, the PlantArray, which is a lysimeter-based system, provides real-time monitoring of the parameters related to whole-plant water relations such as transpiration rate (Tr), growth rate and water use efficiency (WUE)^8^. Genotypic differences on these traits in response to progressive soil stress have been delicately revealed in cowpea and tomato using PlantArray^9-11^.

Legumes are staple foods and important vegetables for many cultures worldwide^12^. Water deficiency at any stage, especially during the grain filling and reproductive phases, can affect legume plant growth and ultimately reduce yield and plant biomass^13^. The yield loss depends on the intensity and duration of drought, crop genotype and developmental stage^14^. Soybean (*Glycine max*. L) native to East Asia is the most important legume crop globally, while cowpea (*Vigna. unguiculata*. L) indigenous to West Africa is one of the most drought-tolerant vegetable legumes popular in Asia^15,16^. The two crops are known to exhibit different regulatory modes of water consumption when suffering drought stress, and have been frequently used in comparative studies of the shoot responses to soil water deficiency^2^. Cowpea typically demonstrates a slower rate of water loss and hence a higher leaf relative water content (RWC) and water potential than soybean as soil water is depleted, which is attributed to the better stomatal control^3,17^. To date, the molecular mechanisms underlying the contrasting water use behaviors between the two species remain largely unknown.

In this study, we used the transpiration-interfaced automatic feedback irrigation function in PlantArray to generate consistent gradual soil drought for cowpea and vegetable soybean (known as maodou) plants. This enabled cross-species physiological and molecular comparisons in the shoot to be made under similar soil drought conditions while the ambient environments, including solar intensity and period, temperature and vapor pressure deficit (VPD), were identical. It also allowed for precise sampling of plant tissues under the intended soil water contents. Our results provide a high-resolution and comprehensive view on the physiological and molecular basis of the profligate versus conservative water use behaviors.

## Results

### Whole-plant water relations under progressive soil water deficit

By deploying feedback irrigation to each pot according to the daily water loss (Table S1, see details in Materials and methods), we generated comparable strengths of gradual soil drought for cowpea and soybean plants and divided the treatment into four periods: the well-watered (WW), moderate soil drought (MD), severe soil drought (SD), and recovery phases (RC, Fig. 1). In both crops, the Tr exhibited a diurnal pattern with the maximum value being observed near noon (Fig. 2A). The WUE, which was expressed as the amount of biomass produced per unit of water used by a plant (g fresh weight g^-1^ water transpired), was greater in cowpea (0.065) than in soybean (0.054) under WW conditions, suggesting more efficient water use in the former (Fig. 2B). A comparison of the daily Tr between the WW and treatment groups showed no significant differences in cowpea during the initial days of irrigation reduction; in contrast, irrigation reduction had an immediate stimulatory effect on Tr in soybean that lasted for six days until the volumetric soil water content (VWC, the volume of water per unit volume of soil) reached a moderately low level of 0.27 (Fig. 2A, C). As drought progressed, Tr significantly decreased in both species. In soybean specifically, significant phase advancement of the midday Tr started to show on the fifth day of treatment (VWC=0.35), and the extent of the phase shift increased with decreasing soil VWC (Fig. 2C). On the seventh day when the drought was severe (VWC=0.18), the maximum phase change (1.9 hours) was observed (Fig. 2C). Following water resumption, the phase difference between the two groups rapidly disappeared. The phase difference of Tr was marginal in cowpeas in the whole course of the experiment.

**Fig. 1.**
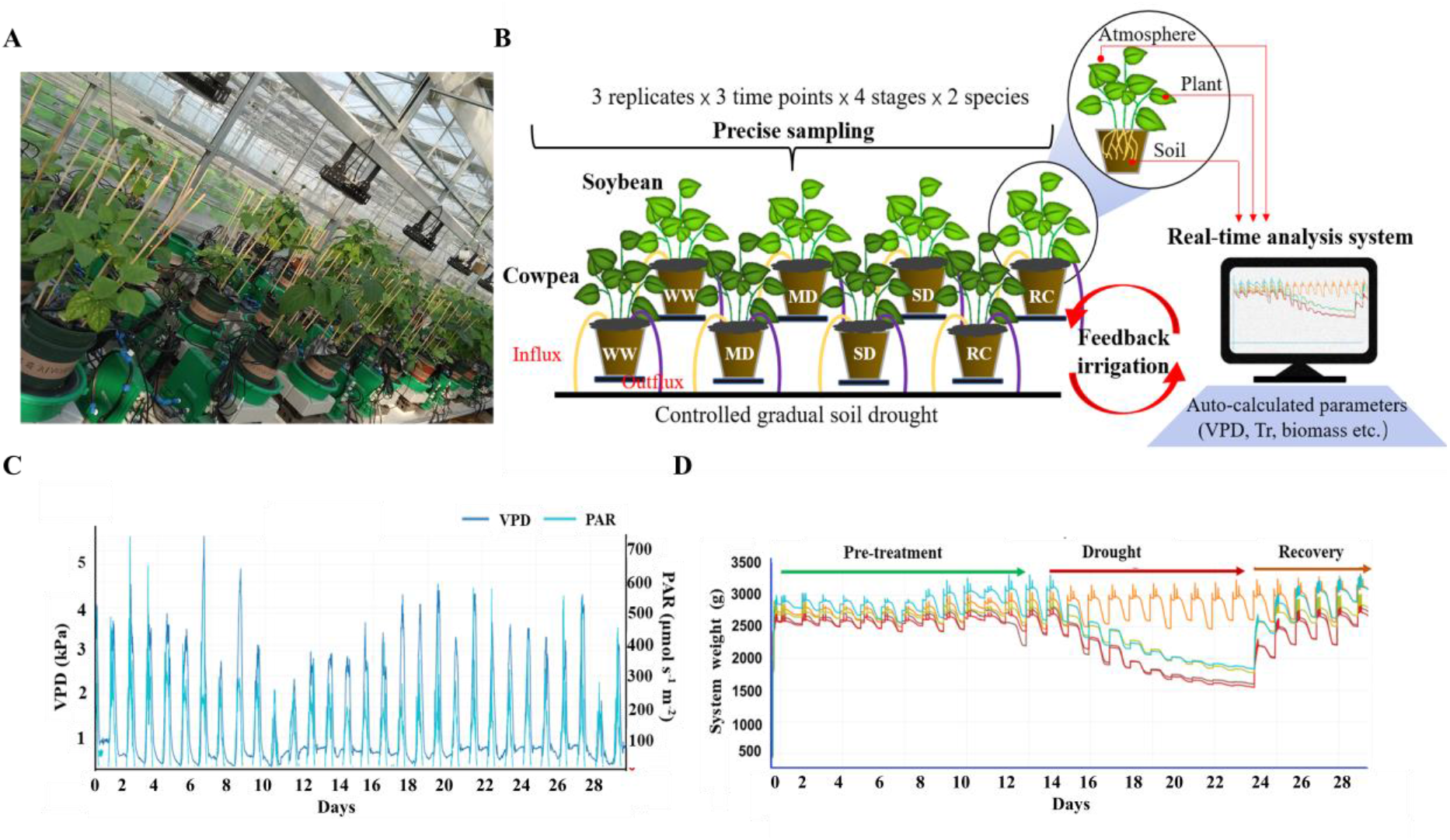
Overall experimental design. A and B, The lysimetric system where randomized experimental array consisting of multiple measuring units loaded with cowpea or soybean plants was set up. PlantArray combines gravimetric system, atmospheric and soil probes, irrigation valves and controller in a unit. A real-time analysis system performs continuous and high-throughput statistical analysis for multiple sensors and sources (atmosphere, plant and soil). The parameters related to whole-plant water relations such as dynamic vapor pressure deficiency (VPD), transpiration rate (Tr) and biomass were subsequently auto-calculated. Three plants grown in each pot and 12 pots of soybean or cowpea were set for phenotyping. Each pot was filled with 3.9-L vermiculite and nutrient soil mixed in a 2:1 (v:v) ratio whose surface was wrapped with plastic film to prevent evaporation, and was irrigated by using an automated feedback system. For sampling, leaf samples were collected from different pots containing cowpea or soybean plants that were under well-watered (WW, Day12), moderate soil drought (MD, Day19), severe soil drought (SD, Day22) or recovery (RC, Day23) treatments, respectively. For each single treatment, one leaflet each from the same trifoliate leaves of the three seedlings in a pot was collected and pooled, at 6 am, 12 pm and 4 pm of the sampling day, respectively. Three biological replicates were analyzed per sample. Totally, 72 RNA-Seq libraries were constructed from leaf samples collected under various soil droughts. C, VPD and photosynthetically active radiation (PAR) conditions during the course of the experiment. D, Dynamics of system weight during the course of the experiment, which consisted of the pre-treatment, drought stress and recovery phases.

**Fig. 2.**
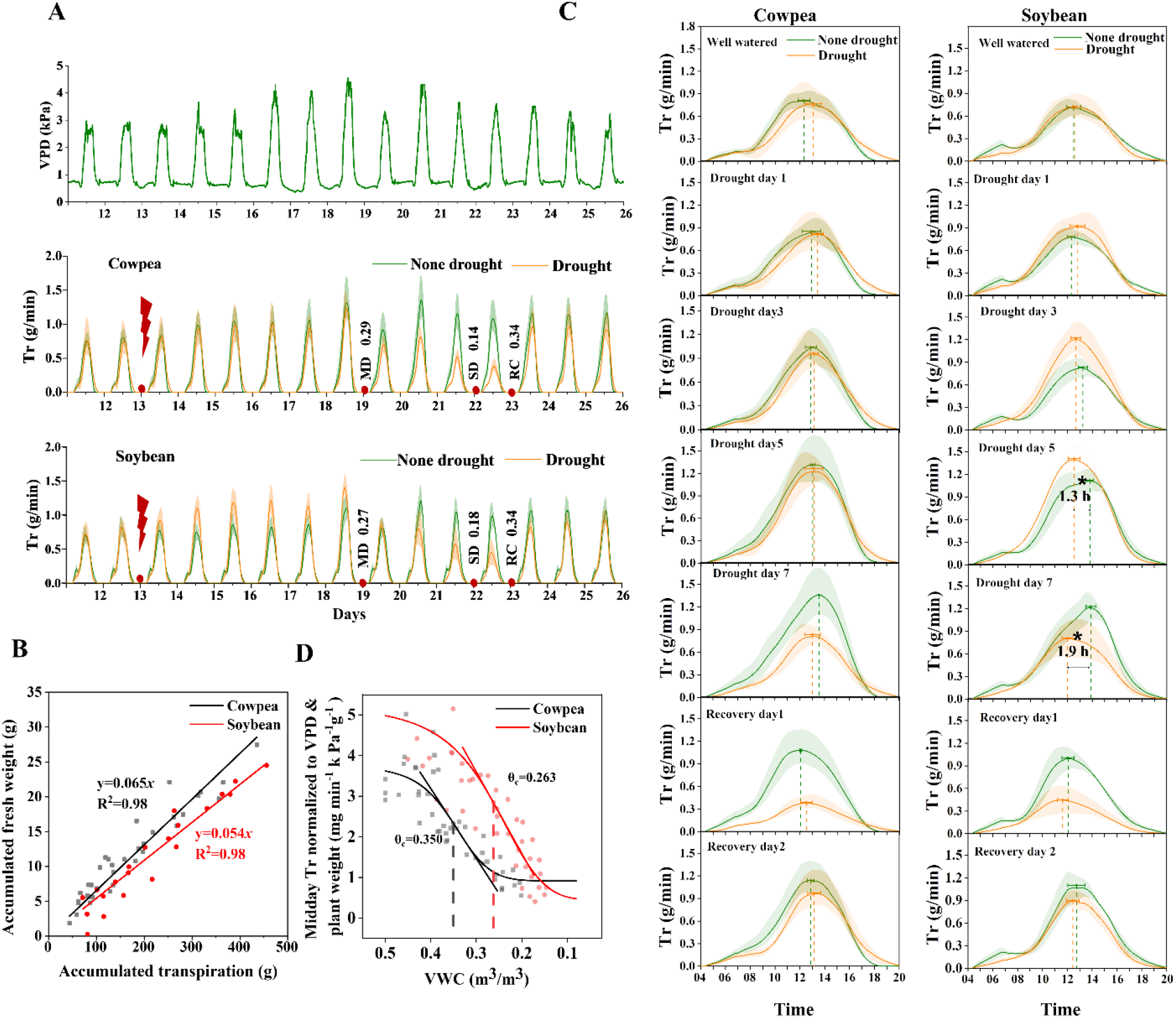
Whole plant water relations of soybean (‘ZN6’) and cowpea (‘TZ30’). A, Dynamics of transpiration rates (Tr) of the well-watered (WW) and drought-stressed cowpea and soybean plants. The date of the onset of irrigation reduction is marked with a flash icon and the VWCs (volumetric moisture content of soil) at the time of sampling in moderate soil drought (MD), severe soil drought (SD) and recovery (RC) stages are shown. B, Water use efficiency (WUE) of the two crops under well-watered condition. WUE was expressed as the amount of biomass produced per unit of water used by a plant. C, Daily Tr of the WW and stressed plants on specific days following treatments. Note the phase shift of midday Tr during MD in soybean. Asterisk indicates significant difference according to a *t*-test at the 5% level of significance. D, Plot of the midday Tr of the drought-stressed cowpea and soybean plants normalized to vapor pressure deficiency (VPD) and plant weight against VWC. θ_c_ represented the inflection point of soil VWC at which a plant showed the fastest decrease of Tr through stomatal control.

Since Tr alone could not exclude the impact of canopy size and the environment, midday Tr (Tr_m_) between 12 pm and 2 pm was then normalized to VPD and plant weight. As shown in Fig. 2D, the daily Tr_m,VPD_ of each crop decreased gradually with the declining VWC, then fell down linearly, and ultimately approached the minimum. To describe the response curves of transpiration against water stress, we used a logistic function to fit Tr_m,VPD_ with VWC, where the maximum and minimum values of Tr_m,VPD_, the maximum decline rate of Tr_m,VPD_ and the corresponding critical VWC (θ_c_) determined the shape of the Tr_m,VPD_-VWC curves. Here, θ_c_ represented the inflection point of soil VWC at which a plant showed the fastest decrease of Tr through stomatal control. The results showed that under WW conditions, the Tr_m,VPD_ in cowpea reached a maximum rate of 3.747±0.421 mg H_2_O min^-1^ kPa^-1^ g^-1^ fresh weight, which was lower than that of soybean (5.247±0.818 mg H_2_O min^-1^ kPa^-1^ g^-1^ fresh weight), representing a more conservative water use strategy (Fig. 2D). The θ_c_ was much greater in cowpea (0.350±0.017 m^3^/m^3^) than in soybean (0.263±0.031 m ^3^/m^3^), indicating that cowpea started constraining water loss under a more abundant soil water condition and corroborating it as a more conservative water consumer. Under very severe soil drought (VWC<0.15), cowpea tended to maintain a higher level of midday Tr_m,VPD_ (0.913±0.279 mg H_2_O min^-1^ kPa^-1^ g^-1^ fresh weight) than soybean (0.451±0.205 mg H_2_O min^-1^ kPa^-1^ g^-1^ fresh weight, Fig. 2D). When irrigation was resumed, the Tr of both species recovered quickly to ∼80% of the WW level within 2 days (Fig 2A).

### Overview of the transcriptomic data

To disclose the gene regulatory basis underlying water use behaviors, RNA-Seq was performed for cowpea and soybean leaves collected at WW, MD, SD and the RC stages. At each sampling stage, the VWC was similar between the crops. A range of 38.2 to 70.6 million high-quality reads were generated from the 72 RNA-Seq libraries constructed from leaf samples collected under various VWCs (Table S2). High mapping rates (> 92.4%) to the reference genomes were observed for all the libraries. Hierarchical cluster analysis showed high correlation among the biological replicates of each sample and a general trend of sample clustering more by Time of day (TOD) than by treatment (Fig. S1). According to the Spearman correlation coefficients (SCC), VuMD-6&VuWW-6 (SCC=0.972) and GmMD-6&GmRC-6 (SCC=0.971) were the most correlated sample pairs in cowpea and soybean, respectively, while the VuMD-6&VuSD-16 (SCC=0.313) and GmRC-6&GmMD-16 (SCC=0.394) pairs were the least correlated. In general, GmMD-16 and VuSD-16 were the samples showing the least correlation to others in the two crops, respectively (Fig. S2A). TOD had a significant impact on both crops and showed complex interactions with the drought scenario (Fig. S2B). For example, the vast majority of cowpea differentially expressed genes (DEGs) were detected at midday under MD but late day under SD. In soybean, the least and most DEGs were recorded in the morning under MD and SD, respectively. A quantitative RT-PCR analysis of randomly selected genes confirmed the accuracy of the RNA-Seq data (Fig. S3).

### Transcriptomic reprogramming as revealed by pairwise comparisons

Pairwise comparisons between the samples collected at the same TOD in different treatment scenarios revealed a total of 4,739 (15.9% of the 29,773 protein-coding genes) and 10,330 (19.5% of the 52,872 protein-coding genes) unique DEGs in cowpea and soybean, respectively (Fig. S2C). The top 20 DEGs were analyzed for each of the nine pairwise comparisons in the two crops (Fig. S4, Table S3). Among them, the ABA signaling gene *HIGHLY ABA-INDUCED PP2C GENE 3* (*HAI3*) and the aquaporin gene *TONOPLAST INTRINSIC PROTEIN 2;3* (*TIP2;3*) drew our attention because they were only found in the top DEG list of cowpeas and were specific to the MD stage, thus were likely related to water conservation in cowpea under such condition. In soybean specifically, multiple *FASCICLIN-LIKE ARABINOGALACTAN* (*FLA*) genes, which are putatively involved in cell expansion and cell wall architecture^18^, were among the top downregulated DEGs.

To more comprehensively understand the functional relevance between all of the DEGs and water use behavior, gene ontology (GO) enrichment analyses were performed. In the leaves of MD-stressed soybean, GO terms related to photosynthesis, cell wall and fatty acid metabolism were enriched by downregulated genes; in contrast, most of these GO terms were not enriched in cowpea leaves suffering from MD, which was consistent with a dehydration avoidance mechanism known for this crop (Fig. S5). As the soil drought became more severe, the two species exhibited more similar GO enrichment profiles, including those related to photosynthesis, cell wall, carbohydrate metabolism and protective/repair processes. Despite these commonalities, we observed interesting species-specific regulations. For example, GO terms related to phosphorus metabolism/phosphorylation and posttranslational protein modification were enriched by upregulated genes only in soybean, while developmental process and aromatic compound biosynthetic process that is related to secondary metabolism were enriched in cowpea (Fig. S5), implying more versatile drought-coping strategies in cowpea.

### MD and SD differentially affected clock gene expressions in the two crops

Since an interaction between TOD and drought scenario was indicated (Fig. S2B), we then examined the dynamic expressions of the clock genes to better elucidate how soil drought at different strengths affected the trend of these genes. A total of 32 and 44 putative circadian clock genes, respectively, were identified from cowpea and soybean (Table S4, Fig. S6). The majority (56 out of the 76) of these genes kept their expression unchanged by drought treatment, while those with changes could be divided into two categories: 1) only the amplitude of gene expression was changed and 2) both the amplitude and daily trend of gene expression were changed (Fig. 3). Four genes in cowpea, including *ONE PSEUDO-RESPONSE REGULATOR 1*(*PRR*), two *EARLY FLOWERING 3* (*ELF3*) and one *ELF4* ortholog, and six soybean genes, including one each of *ELF3, ELF4, ZEITLUPE FAMILY* (*ZTL*), and three *PRR* orthologs, fell into the first category. The second category comprised five genes each in cowpea (two *REVEILLE* (*RVE*), one each of *PRR, ELF4* and *CCA1 HIKING EXPEDITION* (*CHE*) ortholog) and soybean (one each of *ELF3, ELF4, PRR, CHE* and *ZTL* ortholog). Of these 20 genes, all but *GmELF3 like-4* were upregulated by drought, and the *RVE* and *ZTL* orthologs were species-specifically regulated. MD and SD showed different impacts on these genes. For example, while all the *PRR* orthologs of soybean and cowpea were upregulated under SD, only *PRR3* and *PRR5* were responsive to MD. MD and SD had similar impacts on expression trends of the cowpea *ELF4* orthologs, whereas these drought scenarios differentially regulated the expression of the two soybean *ELF4* orthologs, particularly *GmELF4 like-3*. Drought shaped the trends of the expression of *CHE* orthologs in the two species alike despite their different daily patterns under WW, but the amplitude of expression was significantly higher in soybean under SD.

**Fig. 3.**
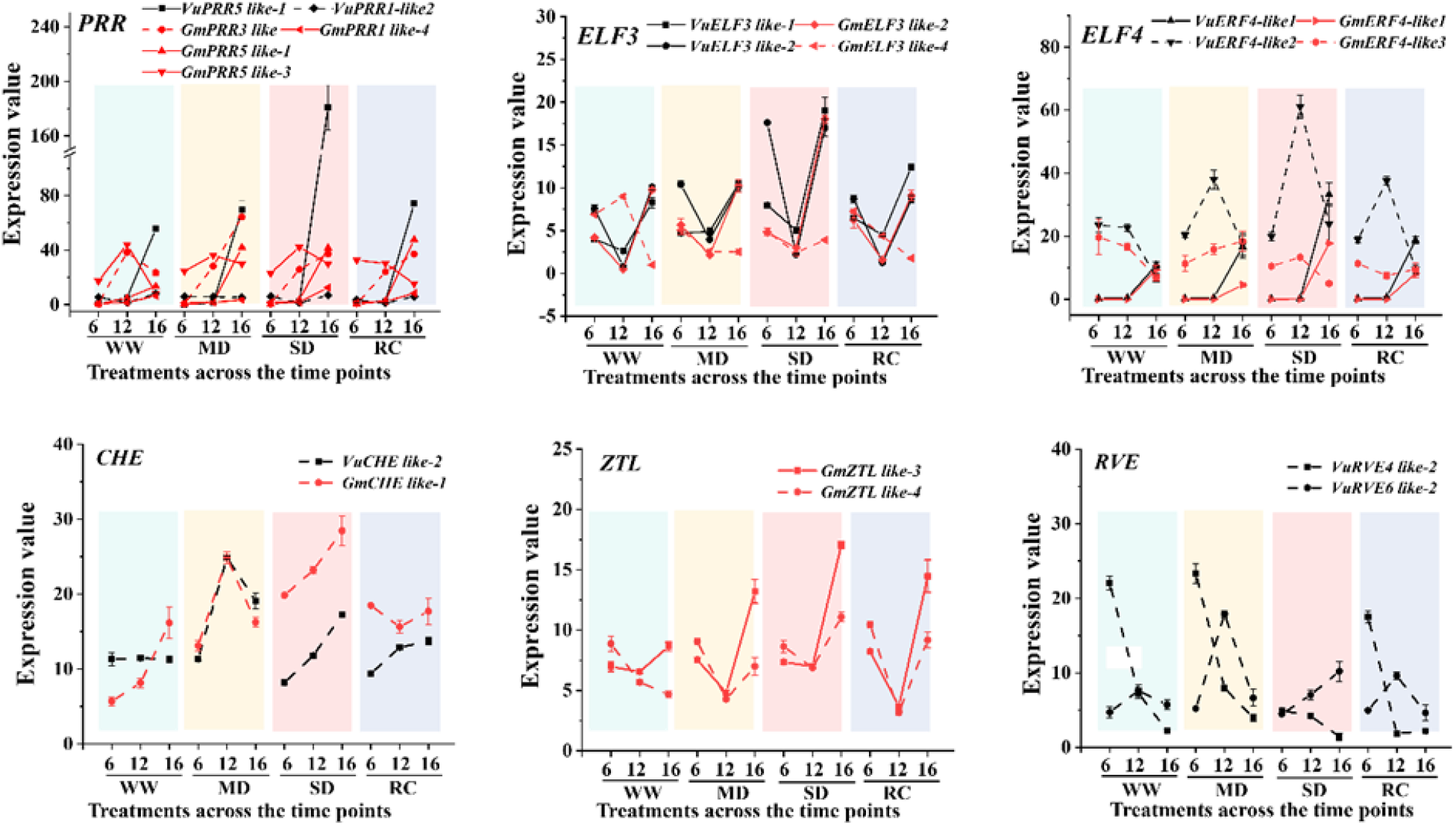
Expressions of the 20 circadian clock genes showing altered amplitude or trend in the two crops. Black line: cowpea; Red line: soybean; Solid lines denote that only the amplitude of gene expression was altered and dotted lines denote that both the amplitude and phase of gene expression was altered. These genes encode afternoon-phased components (*RVE4* and *RVE6*), evening-phased components (*ELF*3, *ELF4, PRR5, PRR1* and *CHE*), and *PRR1*’s regulators, including *PRR3* and *ZTL*. Genes were named according to the Arabidopsis orthologs as shown in Supplemental Table S4, WW: well-watered control; MD: moderate soil drought; SD: severe soil drought, RC: recovery phase.

### Multi-multifactorial gene clustering analysis revealed species-specific gene regulations by drought, TOD and their interactions

To comprehensively identify genes affected by TOD, drought or their interactions, a generalized linear model (GLM)-based analysis of expression was then performed using *EdgeR* implementing custom codes (Data S1). We identified 5,403 soybean genes as affected by drought, 651 by TOD, and 208 by their interactions. In cowpea, the numbers were 1,766, 499 and 30, respectively (Table S5). The aforementioned genes were further clustered based on their expression modes using the k-means method. A total of 28 (12 by drought, 9 by TOD, and 7 by TOD × drought) and 39 (21 by drought, 12 by TOD, and 6 by TOD × drought) clusters were grouped in soybean and cowpea, respectively. Despite the fewer DEGs identified in cowpea as being responsive to drought or TOD × drought, they formed more or similar number of clusters than in soybean, suggesting higher heterogeneity of their expression regulations in cowpea. Functional annotations of each cluster were shown in Table S5.

We were interested in gene clusters showing different expression patterns (*P* ≤ 0.05, Table S6) between MD and SD, because they may harbor the genes that were preferably expressed in either earlier or later stage of drought responses. When considering the significance of GO enrichment (FDR ≤ 0.05), the gene clusters with annotations in cowpea all fell into the drought-responsive category, while those in soybean fell into the TOD- and TOD × drought-responsive categories as well. The cowpea gene clusters were functionally related to cell wall biogenesis, plastid, response to light stimulus, reproductive shoot system development, and ribosome (Fig. S7). In contrast, the soybean gene clusters were involved in oxidoreductase activity, cell cycle, cell periphery, transmembrane transporters, in addition to cell wall. Among them, the GO terms related to ion transport were enriched only in TOD- and TOD × drought-responsive clusters. Clearly, the soil drought scenario and TOD interplayed in different ways in the two crops to affect the shoot gene expressions, which contributed to the formation of the contrasting water use behaviors (see discussions).

### Gene co-expression network and hub genes related to the transpiration rate

Through a weighted gene co-expression network analysis (WGCNA), we identified 17 and 20 gene modules in soybean and cowpea, respectively (Fig. S8, S9A). Two co-expression modules each, viz. modules 9 and 17 (VuM9 and VuM17 hereafter) in cowpea and the modules 4 and 14 (GmM4 and GmM14 hereafter) in soybean, were significantly associated with Tr (*P* < 0.01), a key physiological trait related to water budgeting^19^. GO enrichment analysis revealed common terms in these four modules, such as plastid/chloroplast, responses to radiation and temperature, and responses to stress (Fig. S9B), which are known to relate to stomatal functions, thus corroborating the links of these modules with transpiration. We noted that VuM9 was enriched with many RNA/transcription-related GO terms, suggesting that the water conservation traits in cowpea leaves involved also active gene regulations, rather than was merely a consequence of physiological drought avoidance. On the other hand, GmM14, but not VuM17, was enriched with the GO terms “endomembrane system” and “endoplasmic reticulum”, despite the two modules both were primarily responsive to SD (Fig. S9B, Fig. S10), implying a severely perturbed cellular homeostasis in soybean leaves ^20^. Next, we interlinked the modules between the two organisms by measuring the correlation between their module eigengenes (MEs) using the WGCNA default ‘relating modules to external information’ analysis. This analysis found that, among the Tr-associated modules, the VuM9 was positively correlated with the GmM4 (r=0.721, *P*=0.008), while the VuM17 was negatively correlated with the GmM14 (r=-0.902, *P*=5.89E^-5^). Notably, the GO term “response to heat” fell into the two negatively correlated modules (see discussions).

By using the criteria of intramodular connectivity (*K*_ME_) ≥ 0.85 and absolute gene significance (GS) ≥ 0.85, we identified 5 and 8 hub genes from VuM17 and VuM9, respectively (Fig. 4A, B, Table S7). From GmM4 and GmM14, we identified 26 and 9 hub genes, respectively (Table S7). The soybean hub genes included those encoding various types of transcription factors, heat shock proteins, transporters, and noticeably, the ABA-degrading enzyme abscisic acid 8’-hydroxylase 4 (*CYP707A4*)^21^. The *GmCYP707A4* was transcriptionally upregulated at noon under MD and had a positive GS value, indicating that its expression increased Tr; however, its ortholog in cowpea was downregulated and less abundantly expressed under the same stress despite its higher expression under the WW and SD conditions (Fig. 4C). Given the close relationship between ABA and stomatal closure, the higher expression of *GmCYP707A4* under MD is postulated to relate to the more profligate water use trait in soybean leaves under this specific soil drought scenario. The hub genes of cowpea included those encoding the ABA-signaling component phosphatase 2C family protein, trehalose-phosphatase/synthase 9 (TPS9), RAB GTPase homolog A6B protein, far-red elongated hypocotyl 1 and so on. Among them, *VuTPS9* (*Vigun02g076000*) aroused our particular interest (Fig. 4A), because trehalose has emerged as an important singling molecule in stress responses^22,23^. The expression of *VuTPS9* showed an obvious fluctuation across a day, and was higher under MD and SD than WW. In contrast, the soybean ortholog of *TPS9* (*Glyma*.*02G033500*) showed rather stable expression over the course of gradual drought (Fig. 4C). Intra-modular gene network further showed that *VuTPS9* interacted, at the highest weight, with the genes encoding TPS11, ELF3, the energy-related 3-methylcrotonyl-CoA carboxylase (MCCB), and circadian-related cryptochrome 1 (CRY1) (Fig. 4A, Table S8).

**Fig. 4.**
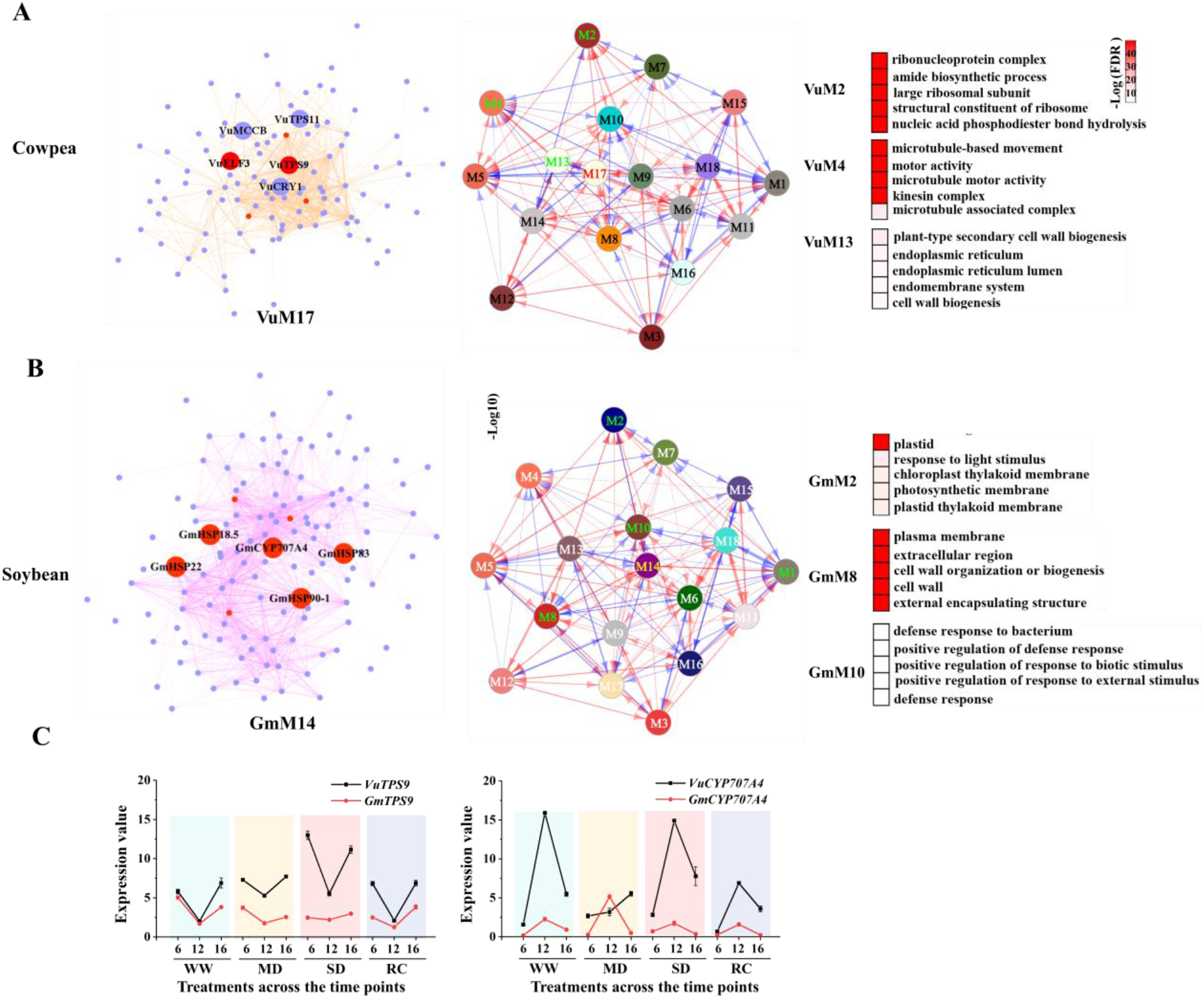
Weighted correlation network analysis. A, Inter-modular network of the gene co-expression modules in cowpea (central panel), intra-modular gene network for the module VuM17 (left panel), and the functional annotations of the most interactive modules (VuM2, VuM4, VuM13) with VuM17. B, inter-modular network of the gene co-expression modules in soybean (central panel), intra-modular gene network for the module GmM14 (left panel), and the functional annotations of the most interactive modules (GmM2, GmM8 and GmM10) with GmM14. Only the top five enriched GO enrichments according to false discovery rate were shown. GmM1, which was also an interactive module with GmM14 and top-enriched with “plastid”, is not included in this figure because of the GO enrichment FDR values beyond the threshold (0.05). Red dots (left panels in A and B) denote hub genes; Medium purple dots (left panel in A and B) denote other genes in the network of module VuM17 or GmM14. Arrowed lines in blue and red color (central panels in A and B) denote positive and negative interactions, respectively. C, Dynamic expression patterns of the *CYP707A4* and *TPS9* orthologous genes in the two species under various drought scenarios. WW: well-watered control; MD: moderate soil drought; SD: severe soil drought, RC: recovery phase. 6, 12, 16 denoted time of day (6 am, 12 pm and 4 pm).

### Functional validation of *VuTPS9* for its role in regulating transpiration rate (Tr)

Among the aforementioned hub genes, *TPS9* and its orthologs had not been functionally characterized in any plant species. Moreover, the cowpea and soybean *TPS9* orthologs exhibited different regulatory patterns under drought treatment, *VuTPS9* was hence selected for functional validation. Gene family analysis identified 10 *TPS* genes from the cowpea genome, and the phylogenetic tree suggested that *VuTPS9* belongs to the Class II subfamily (Fig. S11). Here, 5% and 10% PEG-6000 treatments were adopted to mimic the different strengths of osmotic stress. The Tr and stomatal conductance (Gs) of leaves were compared between the 35S::*VuTPS9-eGFP* (*VuTPS9*-OE) and 35S::*eGFP* overexpression lines, which showed no significant difference before treatment (Fig. 5A, Fig. S12). Treatment with 5% PEG reduced Tr and Gs to a similar level in the two lines on each of the three days during the experiment; 10% PEG treatment caused a rapid and sharper decline in Tr and Gs in both lines, particularly in the *VuTPS9*-OE plants (Fig. 5A, Fig. S12). To better display the daily inhibitory effects in the two lines, the relative Tr and Gs to the 0-day value were calculated. As shown in Fig 5B and Fig. S11C, on the first day of 10% PEG treatment, the levels of Tr and Gs decreased by 92% and 95%, respectively, in the *VuTPS9*-OE plants, as opposed to 57% and 73% in the 35S::*eGFP* line. Consistent with this phenotype, the stomatal aperture in the former line was more sensitive than that in the latter line, as measured 1 day after 10% PEG treatment (Fig. 5C, D).

**Fig. 5.**
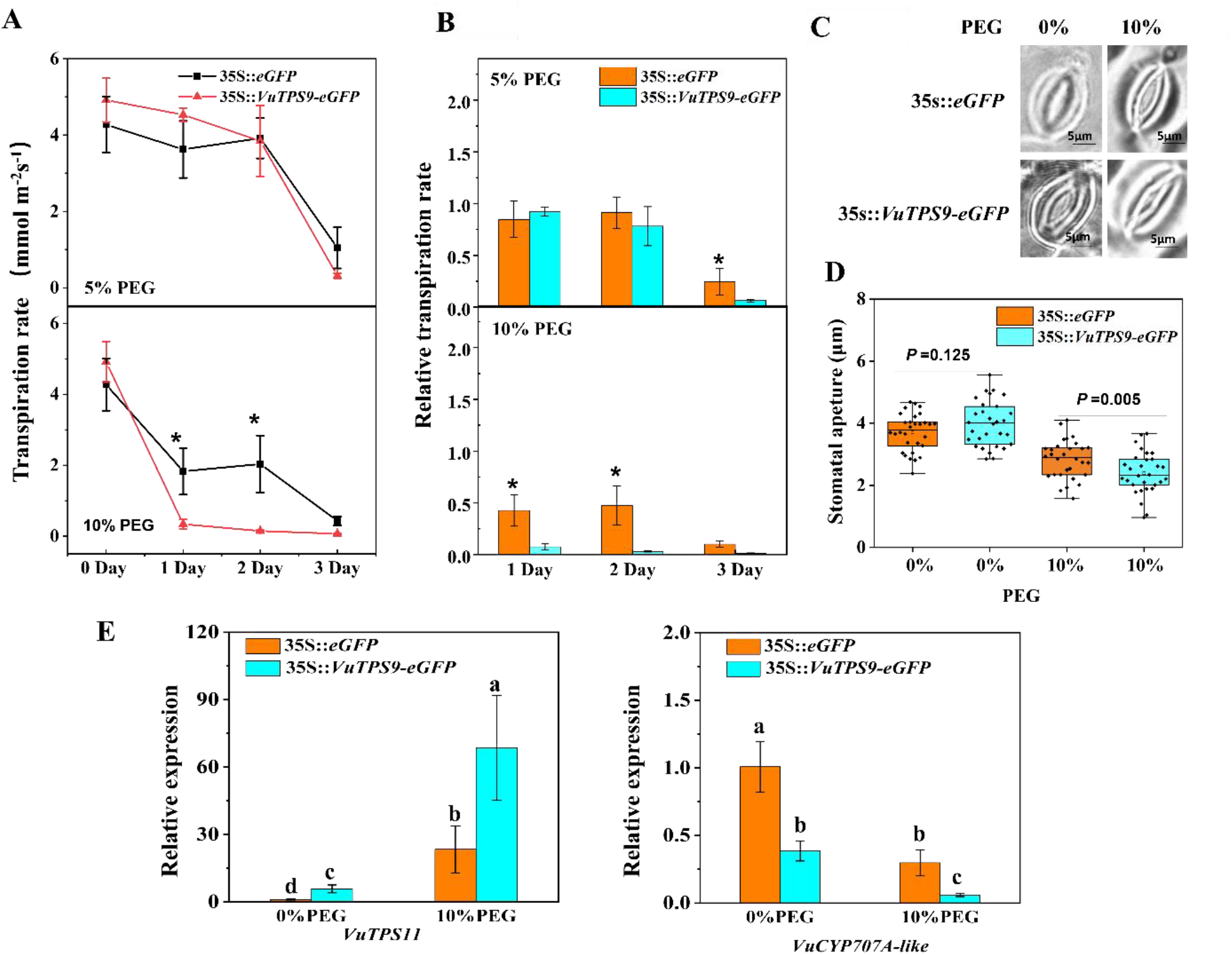
The effect of *VuTPS9* overexpression on transpiration rate and stomatal closure. A and B, The transpiration rates (Tr, A) and relative Tr (B) in the leaves of *VuTPS9*-OE (35S::*VuTPS9-*eGFP) and 35S::*eGFP* lines after PEG-6000 treatment at different concentrations for 0, 1, 2 and 3 days. To calculate the relative Gs, the average Tr amount on day 0 of each line was set as 1. Data are means of at least three biological replicates and are shown with vertical error bars (±SD). Asterisk indicates significant differences according to a *t*-test at the 5% level of significance. C, Representative stomatal images of the two lines exposed to 10% PEG-6000 or mock (water) treatments. D, Stomatal aperture values. Dots represent data measured from 30 stomata under each treatment. *P*-values were calculated from a *t*-test. E, Relative expressions of *VuCYP707A-like* (*Vigun03g186900*) and *VuTPS11* (*Vigun03g378000*) in leaves of *VuTPS9*-OE and 35S::*eGFP* lines after PEG-6000 treatment for 1day. Different letters indicate significant differences according to a Tukey’s test at 5% level.

Lastly, to test the hypothesized co-players of *VuTPS9* based on WGCNA, we examined the expressions of the aforementioned *VuTPS11* (*Vigun03g378000*), which was co-expressed with *VuTPS9* in the module VuM17, as well as *VuCYP707A-like* (*Vigun03g186900*), which was a member of the VuM17-interacting module VuM2. These two genes were selected because they also were putatively involved in trehalose signaling and ABA degrading, respectively. We found that the expressions of both genes were indeed altered in leaves of the *VuTPS9*-OE plants relative to the control line. Expression of *VuTPS11* was enhanced by *VuTPS9* overexpression and 10% PEG treatment, while an opposite trend was observed for *VuCYP707A-like* (Fig. 5E). Therefore, trehalose and ABA signaling appeared to interplay to confer the water use behavior in cowpea.

## Discussion

Transpiration-interfaced automatic feedback irrigation based on soil VWC could accurately control the speed of soil water depletion in each pot, thus creating highly comparable drought scenarios in different pots. This approach reduces the confounding effects of leaf size and growth rate^17^ and maximizes the likelihood of finding genotypic drought response differences. The whole plant transpiration-based assay avoids problems of upscaling conventional leaf-based traits, such as photosynthetic carbon isotope discrimination, to the plant or field scale^24^. The term “physiolomics” was recently proposed to refer to this emerging subject of high-throughput physiology-based phenotyping^25^. Benefiting from the long-term phenotyping and modeling of Tr_m,VPD_, we discovered that cowpea had the potential to maintain a higher transpiration rate under very severe soil drought (VWC < 0.15). This finding is reminiscent of an earlier study reporting lower lethal RWC values of leaves in cowpea (40%) than in soybean (50%)^26^, and argued that conservation or profligation in water use is a conditional concept. Given the viewpoint that any plant traits to avoid or postpone drought become ineffective under terminal drought or in soils with very low water-holding capacity^2^, we propose that the water regime-dependent conservative/profligate water use behavior in a plant is an adaptive trait to help balance survival and productivity under dynamic environmental changes. The high resolution and continuity of the physiolomic assay also led to the detection of the soybean-specific short-term increase along with a phase change of Tr_m_ under MD conditions. Making conclusive interpretation to this phenomenon is still difficult at present, but it might be due to an interplay between MD and midday high temperature. Tr is known to be related to canopy temperature^27^, and soybean is more heat sensitive than cowpea^15,28^. Our assumption could find some support from the molecular data that orthologs of the clock gene *ELF3*, which also is a thermal responsiveness component^29^, showed different regulations by drought between cowpea and soybean. WGCNA further uncovered a negative correlation of two Tr-associated co-expression modules comprising the GO term “response to heat” between the two crops, reinforcing that heat responses may underpin the phase change of Tr_m_.

The soil drought signal perceived by the roots triggers the accumulation of ABA and the initiation of the ABA signaling cascade, which is key to the induction of stomatal closure in shoots^30,31^. The hormonal action of ABA is precisely controlled by its biosynthesis, catabolism and transduction^32^. We found that the ABA signaling repressor gene *HAI3* was among the top-upregulated genes in cowpea only, and the ABA-degrading gene *CYP707A4* was identified as a soybean-specific hub gene associated with Tr. More interestingly, the regulation of both genes showed a drought scenario-dependent feature. We therefore postulate that the more profound and earlier-stage activation of *HAI3* in cowpea contributed to the higher sensitivity of stomatal control and hence drought avoidance under MD, while the MD-specific upregulation of *CYP707A4* in soybean is likely part of the fine-tuning mechanism to maintain higher Gs through ABA degradation. Taken together, the results suggest a species-dependent subtle regulation of ABA signaling via the balance of ABA signal activation and ABA degradation.

Trehalose, a soluble sugar synthesized via a two-step reaction involving trehalose-6-phosphate synthase (TPS), is known for its role in metabolic and osmotic regulation in a variety of organisms. Recently, the signaling role of trehalose has emerged in plants. Arabidopsis *tps1* and *tps5* (Class II) loss-of function mutants showed insensitivity and hypersensitivity to ABA induction of stomatal closure, respectively, suggesting that the two genes exerted contrasting roles in the adjustment of stomatal aperture under stressed conditions^33,34^. Here, we identified *VuTPS9* belonging to the class II subfamily as one of the hub genes and revealed the different responses of the *TPS9* orthologs to progressive drought between cowpea and soybean. We provided evidence that overexpression of *VuTPS9* increased osmotic stress-induced Gs and Tr inhibition under drought conditions. This effect is similar to that reported for *AtTPS1*, a class I *TPS* gene, but contrasting to that of the class II gene *AtTPS5*, arguing that the biological functions of *TPS* genes may not necessarily be related to their subfamily classifications. Our network analysis further suggests that the trehalose signal works with the circadian signal, ABA and energy metabolism processes to regulate transpiration rate in the shoots under soil drought conditions. Put together, these results add to our knowledge on the roles of the *TPS9* gene in drought stress responses.

The circadian clock also plays essential roles in plant development and stress responses. The reciprocal interaction between clock- and drought-responsive genes has been reported. For example, soybean orthologs of the morning loop component CCA1/LHY were found to negatively control drought tolerance by regulating the ABA pathway^35^. A prevailing transcriptomic reconfiguration by drought stress was observed in the late day in *poplus*^36^. Our results revealed different sensitivities as well as patterns of amplitude and phase of many clock genes to soil drought between cowpea and soybean. Similarly, Li et al. (2019)^37^ reported that the interfaces between the soybean clock and abiotic stress signals were quite different from those in Arabidopsis, suggesting that species-specific phase changes in response to environmental cues are a norm. The phase change of gene expression can have a profound effect on related biological processes, as demonstrated in the case of iron utilization efficiency in soybean^37^. In this study, the expressions of two RVE orthologs in cowpea (but not soybean) showed significant phase changes. These results imply that RVEs may be important in water conservation in legumes. Furthermore, prevailing transcriptomic reconfiguration was observed in the late afternoon, which, however, was more evident at the MD stage for soybean and SD stage for cowpea, respectively. The elevated expression of clock genes in late days has been related to increased ABA synthesis in barley^38^. Given the conserved relationship between ABA and drought response in plants, we assume that the interplay among crop type, drought scenario and TOD had an impact on ABA accumulation and was partly accountable to the contrasting water use behaviors in the two crops. Recently, the influences of the circadian clock on the long-term WUE of Arabidopsis were also reported^39^, adding the importance of this interplay in plant water budgeting.

Collectively, a model elucidating the differential water use strategies in cowpea and soybean is developed (Fig. 6), in which the importance of genes related to water use behaviors such as *VuHAI3, VuTIP2;3, VuTPS9, GmCYP707A4, GmFLAs*, and their putative interactive genes such as *VuELF3, VuCRY1, VuTPS11*, are highlighted.

**Fig. 6.**
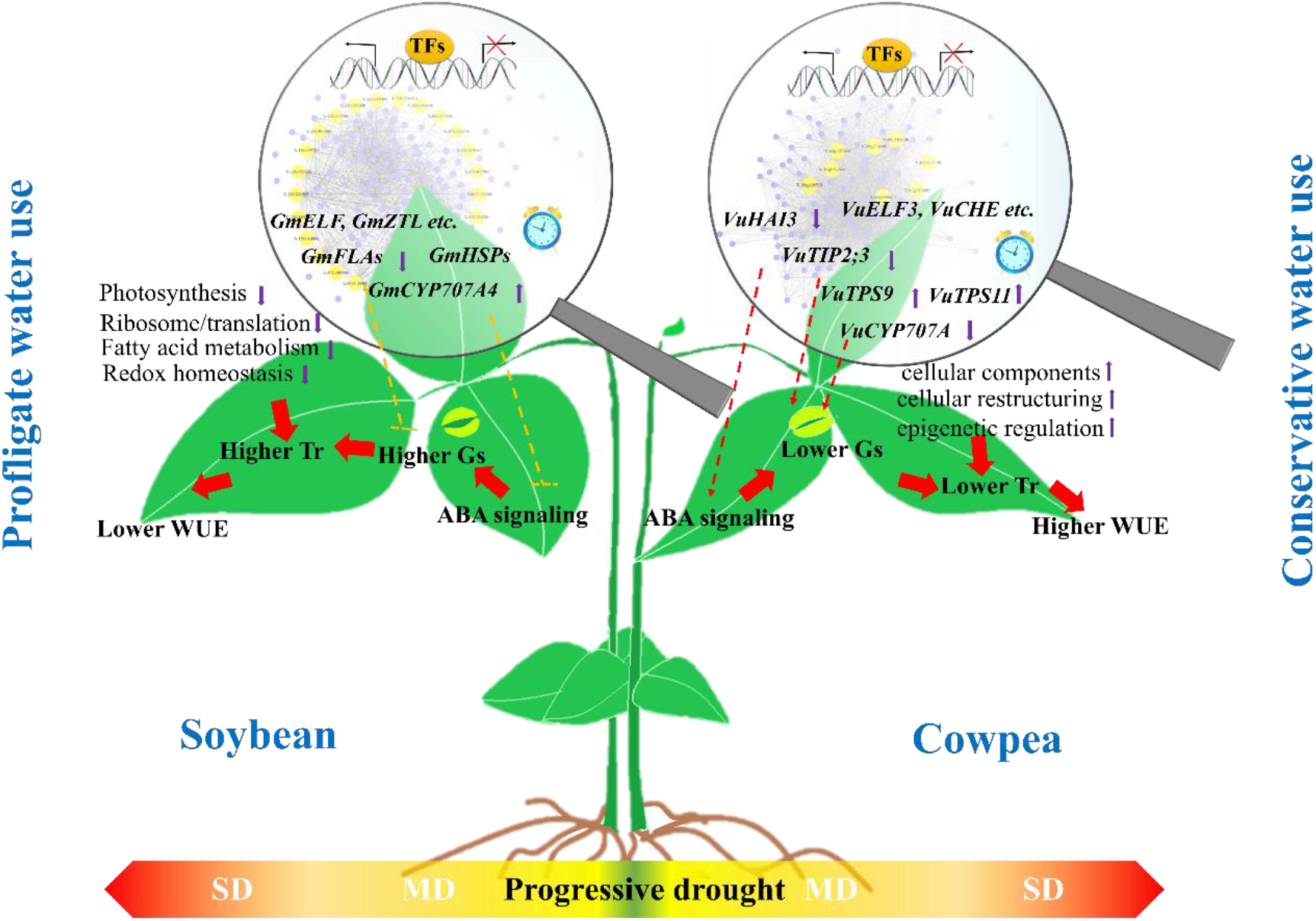
A proposed model of mechanism underpinning the contrasting water use strategies in soybean and cowpea. In cowpea, *VuHAI3*, a negative regulator of ABA signaling was down-regulated under moderate soil drought, thus may trigger the ABA signaling. The down-regulated tonoplast aquaporin gene *VuTIP2;3* and up-regulated trehalose-synthase gene *VuTPS9* are proposed to accelerate the drought-induced stomatal closure. In addition, changes of gene expression patterns of *VuTPS11, VuCYP707A*, are suggested to be involved in the process. Up-regulated genes related to cellular components may modulate water use efficiency via cellular re-structuring and epigenetic regulation. The higher Tr in soybean may be caused by the greater Gs that is related to the high expression of the ABA degradation enzyme *GmCYP707A4*. Besides, photosynthesis, ribosome/translation, fatty acid metabolism and redox homeostasis are quickly destroyed even under MD condition. What’s more, species-specifically regulated circadian clock gene that shape the rhythmic patterns in response to drought may also play a key role in modulating water use strategies. Arrowheads denote the effect of activation and blunt ends denote the effect of inhibition. Up and down arrows indicate up- and down-regulations, respectively. Dotted lines indicate that confirmation is required. TFs: transcription factors; MD: moderate soil drought; SD: severe soil drought; Gs: stomatal conductance; Tr: transpiration rate.

## Materials and methods

### Plant materials and irrigation conditions

Plant materials include one cowpea (‘TZ30’) and one soybean (‘ZN6’) genotype, respectively. The physiological experiment was conducted using the “PlantArray” (Plant-DiTech, Israel) phenotyping platform that installed in a greenhouse in October to November 2019 in Huai’an (33.62^°^N, 119.02^°^E), China (Fig. 1). PlantArray combines gravimetric system, atmospheric and soil probes, irrigation valves and controller in a unit^8^. Four-week-old plants were transferred into the load cells on the system, with three plants grown in each pot and 12 pots set for each species. Each pot was filled with 3.9-L vermiculite and nutrient soil mixed in a 2:1 (v:v) ratio whose surface was wrapped with plastic film to prevent evaporation, and was irrigated by multi-outlet dripper assemblies (Netafim, Israel) that were pushed into the soil and connected to an automated feedback system as described previously^8,10^. Before the start of the experiment, all units were calibrated for reading accuracy and drift levels under constant load weights. The experiment lasted for 28 days and the drought treatment started on day 13, during which gradual deficit irrigation was given during night by the feedback irrigation system that reduced the irrigation levels every day for each pot based on the daily water loss (Table S1). This approach enabled dynamically comparable soil drought strength (feedback drought slope) to be imposed between the two species over time, despite their heterogenous growth rates and plant sizes. Note that measuring leaf area is no longer necessary here with PlantArray for normalized (to biomass) plants^8^. Physiological profiles of the plants in each pot were compiled automatically during the whole experiment which was divided into four treatment periods: WW, MD, SD, and RC. The soil-plant-atmosphere parameters including VPD, photosynthetically active radiation and temperature were monitored simultaneously and continuously with a high resolution (every 3 minutes).

### Data acquisition and analysis of the water relations parameters

Data acquisition was previously described in details by Halperin et al. (2017)^8^. Briefly, the daily whole-plant transpiration was calculated by the difference of measured system weight at pre-dawn and evening. The daily cumulative biomass gain and transpiration throughout a 7-day well-irrigated period was fitted by a linear function, where the slope was determined as the WUE. The plant weight was calculated as the sum of the multiplication of the cumulative transpiration during the period by the WUE and the initial plant weight. The latter, determined as the difference between the total system weight and the sum of the tare weight of pot plus drainage container, weight of soil at pot capacity, and weight of water in the drainage container at the end of the free drainage.

The momentary whole-plant Tr was computed by multiplying the first derivative of the measured system weight by −1. The average whole-plant Tr at mid-day normalized to VPD (Tr_m,VPD_, between 12 am and 4 pm) and the corresponding volumetric moisture content of soil (VWC) were fitted well with a 4-parameter logistic function as the following,

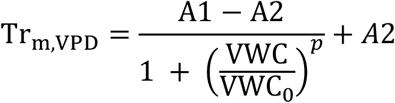

where A1 and A2 were the maximum and minimum values of Tr_m,VPD_ depending on the fitting result at the well-irrigation period and severe-drought period, respectively. *p* indicates the maximum slop of the curve at the inflection point, ie., VWC_0_, where *p* and VWC_0_ mainly determines the sensitivity of Tr_m,VPD_ in response to VWC.

### RNA isolation and RNA-Seq

RNA was extracted using Trizol reagent (Invitrogen, Carlsbad, CA). Five micrograms of total RNA from each sample were used to construct an RNA-Seq library by using the TruSeq RNA Sample Preparation Kit according to the manufacturer’s instructions (Illumina, San Diego, CA). A total of 72 libraries (Table S2) were constructed and then sequenced on the Novaseq 6000 platform.

### Raw data processing and pairwise differential expression analysis

The raw RNA-Seq reads were filtered and trimmed using SeqPrep (https://github.com/jstjohn/SeqPrep) and Sickle (https://github.com/najoshi/sickle) with default parameters. The cleaned reads were combined and aligned to the soybean reference genome WM82.a4.v1 (Phytozome v13) or the cowpea reference genome *Vigna unguiculata* v1.1^40^ using TopHat (V2.1.0). The count of the mapped reads from each sample was derived and normalized to fragments per kilobase of transcript length per million mapped reads (FPKM) for each predicted transcript using Cufflinks (v2.2.1).

Pairwise comparisons were made between samples collected at the same timepoint for each crop. To reduce noises, only genes having an FPKM ≥ 1 in the three replicates of one or both samples in a comparison and a coefficient of variant (CV) < 0.2 among the 3 replicates were considered. The genes exhibiting a difference of at least twofold change with the false discovery rate (*q*-value) ≤ 0.05 were considered as DEGs.

### GLM-based differential expression analysis and clustering of the DEGs

Let y_*i*_ = (y_*e*_(1), y_*i2*_(2), …, y_*ik*_(*t*)) denote the expression measure of gene *i* at different time points of day under treatment *k*. Thus, the GLM that determines the gene expression of gene *i* is expressed as follows:

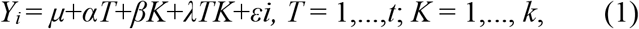

where *μ* is the mean value; *T* is the time vector; *K* is the indicator variables that describe the different treatments; *α* and *β* are the coefficients of time and treatment; *λ* reflects the interaction between time and treatment; and *εi* is the random error, which is generated from a negative binomial distribution. Here, we implemented model (1) for DEGs screening by *edgeR* package in the R platform^41^. We used three-hypothesis tests to detect genes that were differentially expressed in response to the different factors including drought treatment, TOD, and the drought × TOD interaction. An FDR value of 0.05 was adopted as the threshold for significance.

The k-means method was used to cluster the DEGs into distinct groups. Here, the |log_2_FC| value was used as an additional standard to screen the DEGs for higher stringency. The |log_2_FC| threshold was set to 1, 0.5, and 0.25 for the drought, TOD, and drought × TOD category, respectively, by considering the nature of log_2_FC calculation in the model. We applied the between-cluster sum of squares to determine an optimal number of gene groups, using 0.75 as the threshold value. The *t*-test was used to identify whether a significant difference existed between a pair of clusters from different combinations.

### Gene ontology (GO) enrichment analysis

GO enrichments were analyzed for the DEGs using AgriGO V2.0^42^ under a *q*-value threshold of 0.01 for statistical significance. The cowpea reference (787,759 GO assignments for 14,079 genes, 56 assignments/gene) was comparable to that of soybean (1,444,608 GO assignments for 25,073 genes, 58 assignments/gene) in AgriGO V2.0.

### Circadian clock gene analysis

To identify putative circadian clock genes from soybean and cowpea, the sequences of 14 known Arabidopsis core clock proteins^43^ were used as queries in BLAST searches against the soybean and cowpea genomes (Table S4). An *e*-value cutoff of 1*e*^-20^ was applied and the most similar sequence for each BLAST search was selected.

### Weighted gene co-expression network analysis

The co-expression gene network modules were inferred from DEGs using a WGCNA implemented in R. FPKM values of the three replicates were averaged and log transformed. The automatic one-step network construction and module detection method with default settings were used, which include an unsigned type of topological overlap matrix (TOM), a power *β* of 10, a minimal module size of 30, and a branch merge cut height of 0.25. Modules were determined by the dynamic tree cut method. The module eigengene (ME) value, which is defined as the first principal component of a gene module^44^, was calculated and used to test the association of the modules with drought treatment. Spearman correlation between each consensus ME and Tr was calculated to find modules of interest. Lasso regression analysis implemented in the R package “glmnet” was used to construct the inter-modular network through a custom script (Data 1).

### Quantitative RT-PCR

qPCR was performed on qTOWER^®^3 Real-Time PCR detection system (Analytik Jena, Germany) using TOROGGreen^®^ qPCR Master Mix (Toroivd, Shaihai, China). Quantification of relative expression level was achieved by normalization against the transcripts of house-keeping genes^45^ *VuACTIN* and *GmACTIN*. The primer sequences are listed in Table S10.

### Analysis of *VuTPS* genes

The complete protein sequences of TPS of Arabidopsis, rice and soybean were downloaded from Phytozome (https://phytozome-next.jgi.doe.gov/). They were then subjected to BLASTP searches against the cowpea genome (version 1.2). The putative TPS proteins were further confirmed with the conserved domain search tool SMART. In total, 10 putative TPS genes were identified in cowpea. The phylogeny was constructed by using the maximum-likelihood method in MEGA6.0 with 1000 bootstrap replicates.

### Transient overexpression assay

The full cDNA fragment of *VuTPS9* amplified by the primers (F:5’-CGACGACAAGACCGTCACCATGGCGTCAGGATCATATGC-3’; R:5’-GAGGAGAAGAGCCGTCGAGCCATGCTTTCAAAGGAAACT −3’) was cloned into the pVC vector fused with the eGFP sequence at the 3’-terminus. The resulting CaMV 35S::*VuTPS9-eGFP* construct was introduced into *Agrobacterium rhizogenes* strain *GV3101*. Agrobacterium culture containing the construct was resuspended in infiltration buffer (10 mM MES-KOH, pH 5.6, 10 mM MgCl_2_ and 100 μM acetosyringone) and adjusted to a concentration of OD_600_=1.2. The suspensions were infiltrated into the fully developed unifoliate leaves of 10-day old cowpeas grown in Hogland solution. After one day of transformation, the seedlings were exposed to Hogland solution supplemented with 5% or 10% PEG-6000 for 3 days. The expression of *VuTPS9-eGFP* was determined by the observation of eGFP signals.

### Portable device-based measurement of stomatal conductance (Gs) and transpiration rate (Tr)

The Gs and Tr of the transiently transformed leaves were measured using an infrared gas analyzer-based portable photosynthesis system (LI-6400; Li-Cor, Lincoln, NE, USA) mounted with a red/blue LED light source. The environmental condition was maintained at a temperature of 28℃, humidity of 85% and the PPFD of 1000 μmol m^-2^s^-1^. The measurements were made at midday (12 pm to 1 pm) on each day after PEG treatment. Data were acquired from five biological replicates of the control and *VuTPS9-OE* plants.

## Data availability

The RNA-seq raw data are accessible in GenBank with the accession numbers stored in Table S11. The R codes for the GLM, clustering and inter-modular network analyses are available in Data S1.

## Supplemental data

**Table S1**. The irrigation settings for the two species.

**Table S2**. Summary of the sequence data generated from 72 RNA libraries through Illumina sequencing.

**Table S3**. Top 10 up- and downregulated genes in each comparison.

**Table S4**. Orthologs of core circadian clock genes and their expressions in cowpea and soybean.

**Table S5**. Components and functional annotation of the gene clusters that were affected by drought, TOD, or by their interactions.

**Table S6**. *P*-values of clusters that affected by drought, TOD or their interactions according to T-test between MD and SD.

**Table S7**. Hub genes identified from the Tr-associated modules of soybean and cowpea.

**Table S8**. The genes that interacted with VuTPS9 in the module of VuM17.

**Table S9**. Regression coefficients of the gene co-expression modules in soybean and cowpea.

**Table S10**. Primer sequences used for qRT-PCR analysis

**Table S11**. NCBI RSA accession number of the RNA-seq raw data.

**Fig. S1**. Hierarchical clustering of the RNA-Seq data.

**Fig. S2**. Overview of the transcriptomic data.

**Fig. S3**. Validation of RNA-Seq by qRT-PCR on 10 randomly selected genes from the two crops.

**Fig. S4**. Top up- and down-regulated genes in the nine comparisons and their topological relationships.

**Fig. S5**. Gene ontology enrichment analysis of the DEGs under moderate and severe soil drought conditions in comparison to the well-watered condition.

**Fig. S6**. Phylogenetic tree of the orthologs of core circadian clock genes from *Arabidopsis thaliana, Glycine max* and *Vigna unguiculata*.

**Fig. S7**. Gene clusters and their functional annotations in the two crops.

**Fig. S8**. Hierarchical cluster trees showing the modules of coexpressed genes.

**Fig. S9**. Modules associated with transpiration rate in cowpea and soybean.

**Fig. S10**. Eigengene expression of VuM17 in cowpea and GmM14 in soybean.

**Fig. S11**. Phylogenetic tree of the TPS proteins from *Arabidopsis thaliana, Oryza sativa, Glycine max* and *Vigna unguiculata*.

**Fig. S12**. The effect of *VuTPS9* overexpression on stomatal conductance.

**Data S1**. Custom codes used in generalized linear model-based analysis of gene expression, clustering and inter-modular networking.

## Acknowledgment

We thank Wenzhao Xu and Dexu Luo for assistances in the greenhouse experiment, Liang Zeng for assistances in bioinformatical analyses and Menachem Moshelion and Amir Mayo for assistances in running PlantArray. This work is supported by the National Key Research & Development Program of China (Grant No. 2021YFE19800), National Natural Science Foundation of China (Grant No. 31772299, 31861143044), and the Natural Science Foundation of Zhejiang Province (Grant No. LQ21C150004).

## Author contributions

P.X. designed the research and wrote the manuscript. P.F. participated in the experiments, data analysis and wrote the manuscript. T.S. and P.A.K. participated in high-throughput physiological phenotyping analysis. P.F., L.J. and C.S. performed the transcriptomic data analysis. X.W. and Y.H. performed the transient overexpression assay. X.L. participated in the measurement of Tr and Gs.

## References

1. Huber, A.E., Melcher, P.J., Piñeros, M.A., Setter, T.L. & Bauerle, T.L. Signal coordination before, during and after stomatal closure in response to drought stress. New Phytol 224, 675–688 (2019).

2. Blessing, C.H., Mariette, A., Kaloki, P. & Bramley, H. Profligate and conservative: water use strategies in grain legumes. J Exp Bot 69, 349–369 (2018).

3. Likoswe, A.A. & Lawn, R.J. Response to terminal water deficit stress of cowpea, pigeonpea, and soybean in pure stand and in competition. Aust J Agr Res 59, 27–37 (2008).

4. Moroke, T.S., Schwartz, R.C., Brown, K.W., & Juo, A.S. Water use efficiency of dryland cowpea, sorghum and sunflower under reduced tillage. Soil Till Res 112, 76–84 (2011).

5. Klein, T. & Niu, S. The variability of stomatal sensitivity to leaf water potential across tree species indicates a continuum between isohydric and anisohydric behaviours. Funct Ecol 28, 1313–1320 (2014).

6. Vijayaraghavareddy, P. et al. Acquired traits contribute more to drought tolerance in wheat than in rice. Plant Phenomics 2, 16 (2020).

7. Vadez, V. & Ratnakumar, P. High transpiration efficiency increases pod yield under intermittent drought in dry and hot atmospheric conditions but less so under wetter and cooler conditions in groundnut (Arachis hypogaea (L.)). Fielded Crops Res 193, 16–23 (2016).

8. Halperin, O., Gebremedhin, A., Wallach, R. & Moshelion, M. High-throughput physiological phenotyping and screening system for the characterization of plant-environment interactions. Plant J 89, 839–850 (2017).

9. Xu P., Moshelion M., Wu X.H., Halperin O., Wang B.G., Luo J., Wallach R., Wu X.Y., Lu Z.F. & Li G.J. Natural variation and gene regulatory basis for the responses of asparagus beans to soil drought. Frontiers In Plant Science 6, 891 (2015)

10. Pandey, A. K. et al. Functional physiological phenotyping with functional mapping: A general framework to bridge the phenotype-genotype gap in plant physiology. iScience 24, 102846 (2021).

11. Wu, X. et al. Unraveling the genetic architecture of two complex, stomata-related drought-responsive traits by high-throughput physiological phenotyping and GWAS in Cowpea (Vigna. Unguiculata L. Walp). Front Genet 12, 743758 (2021).

12. Rubiales, D., & Mikić, A. Introduction: Legumes in Sustainable Agriculture. Crit Rev Plant Sci 34, 2–3 (2015).

13. Rubiales, D. et al. Achievements and Challenges in Legume Breeding for Pest and Disease Resistance. Crit Rev Plant Sci 34, 195–236 (2015).

14. Zhu, J. Salt and drought stress signal transduction in plants. Annu Rev Plant Biol 53, 247–273 (2002).

15. Lee G., Crawford G.w., Liu L., Sasaki Y. & Chen X. Archaeological Soybean (Glycine max) in East Asia: Does Size Matter? Plos one 6, e26720 (2011).

16. Da Silva, A.C. et al. Cowpea: a strategic legume species for food security and health. London, UK: IntechOpen (2019).

17. Marrou, H., Vadez, V. & Sinclair, T.R. Plant survival of drought during establishment: an interspecific comparison of five grain legumes. Crop Sci 55, 1264–1273 (2015).

18. MacMillan, C.P., Mansfield, S.D., Stachurski, Z.H., Evans, R. & Southerton, S.G. Fasciclin-like arabinogalactan proteins: specialization for stem biomechanics and cell wall architecture in Arabidopsis and Eucalyptus. Plant J 62, 689–703 (2010).

19. Gosa, S.C., Lupo, Y. & Moshelion, M. Quantitative and comparative analysis of whole-plant performance for functional physiological traits phenotyping: New tools to support pre-breeding and plant stress physiology studies. Plant Sci 282, 49–59 (2019).

20. Wang, X. et al. The roles of endomembrane trafficking in plant abiotic stress responses. J Intergr Plant Biol 62, 55–69 (2019).

21. Saito, S. et al. Arabidopsis CYP707As encode (+)-abscisic acid 8’-hydroxylase, a key enzyme in the oxidative catabolism of abscisic acid. Plant Physiol 134, 1439–1449 (2004).

22. Leyman, B., Dijck, P.V. & Thevelein, J.M. An unexpected plethora of trehalose biosynthesis genes in Arabidopsis thaliana. Trends Plant Sci 6, 510–513 (2001)

23. Vishal, B., Krishnamurthy, P., Ramamoorthy, R. & Kumar, P.P. OsTPS8 controls yield-related traits and confers salt stress tolerance in rice by enhancing suberin deposition. New Phytol 221, 1369–1386 (2019).

24. Konate, N.M., Dreyer, E. & Epron, D. Differences in carbon isotope discrimination and whole-plant transpiration efficiency among nine Australian and Sahelian Acacia species. Ann Forest Sci 73, 995–1003 (2016).

25. Li, Y. et al. High-Throughput physiology-based stress response phenotyping: Advantages, applications and prospective in horticultural plants. Hortic Plant J 7, 181–187 (2021).

26. Sinclair, T.R. & Ludlow, M.M. Influence of soil water supply on the plant water balance of four tropical grain legumes. Aust J Plant Physiol 13, 329–341 (1986).

27. Yang G. et al. Unmanned aerial vehicle remote sensing for field-based crop phenotyping: current status and perspectives. Front Plant Sci 8, 1111 (2017).

28. Nahar, K., Hasanuzzaman, M. & Fujita, M. Heat stress responses and thermotolerance in soybean. In: Mohammad M, eds. Abiotic and Biotic Stresses in Soybean Production, UK: Academic Press 261–284 (2016).

29. Zhang, L., Luo, A., Davis, S.J. & Liu, J. Timing to grow: role of clock in thermomorphogensis. Trends Plant Sci 26, 1248–1257 (2021).

30. Sussmilch, F.C., Brodribb, T.J. & McAdam, S.A. What are the evolutionary origins of stomatal responses to abscisic acid in land plants? J Integr Plant Biol 59, 240–260 (2017).

31. Hsu, P.K., Dubeaux, G., Takahashi, Y. & Schroeder, J.I. Signaling mechanisms in abscisic acid-mediated stomatal closure. Plant J 105, 307–321 (2021).

32. McAdam, S. & Sussmilch, F.C. The evolving role of abscisic acid in cell function and plant development over geological time. Semin Cell Dev Biol 109, 39–45 (2021).

33. Gómez L.D., Gilday A., Feil., R., Lunn J.E. & Graham I.A. AtTPS1-mediated trehalose 6-phosphate synthesis is essential for embryogenic and vegetative growth and responsiveness to ABA in germinating seeds and stomatal guard cells. The plant Journal 64, 1–13 (2010).

34. Tian, L. et al. The trehalose-6-phosphate synthase TPS5 negatively regulates ABA signaling in Arabidopsis thaliana. Plant Cell Rep 38, 869–882 (2019).

35. Wang, K. et al. Two homologous lhy pairs negatively control soybean drought tolerance by repressing the aba responses. New Phytol 229, 2660–2675 (2020).

36. Wilkins, O., Waldron, L., Nahal, H., Provart, N.J. & Campbell, M.M. Genotype and time of day shape the populus drought response. Plant J 60, 703–715 (2009).

37. Li, M. et al. Comprehensive mapping of abiotic stress inputs into the soybean circadian clock. P Natl Acad Sci USA 116, 23840–23849 (2019).

38. Habte, E., Müller, L.M., Shtaya, M.J., Davis, S.J. & von Korff, M. Osmotic stress at the barley root affects expression of circadian clock genes in the shoot. Plant Cell Environ 37, 1321–1327 (2014).

39. Simon, N., Graham, C. A., Comben, N. E., Hetherington, A. M. & Dodd, A. N. The Circadian clock influences the long-term water use efficiency of Arabidopsis. Plant Physiol 183, 317–330 (2020).

40. Lonardi, S. et al. The genome of cowpea (Vigna unguiculata [L.] Walp.) Plant J 98: 767–782 (2019).

41. Robinson, M.D., McCarthy, D.J. & Smyth, G.K. edgeR: a Bioconductor package for differential expression analysis of digital gene expression data. Bioinformatics 26, 139–140 (2010).

42. Tian, T. et al. agriGO v2.0: a GO analysis toolkit for the agricultural community, 2017 update. Nucleic Acids Res 45, W122–W129 (2017).

43. Hsu, P.Y. & Harmer, S.L. Wheels within wheels: the plant circadian system. Trends Plant Sci 19, 240–249 (2014).

44. Zhang, B. & Horvath, S. A general framework for weighted gene co-expression network analysis. Stat Appl Genet Mol 4, 17 (2005).

45. Livak, K.J. & Schmittgen, T.D. Analysis of relative gene expression data using real-time quantitative PCR and the 2− ΔΔCT method. Methods 25, 402–408 (2001).

