## Supplemental Table 10 for "Understanding water conservation vs. profligation traits in vegetable legumes through a physio-transcriptomic-functional approach"

**Supplemental Table S****10. Primer sequences used for qRT-PCR analysis**

| **Accession number** | **Gene name** | **Forward primer** | **Reverse Primer** |
| --- | --- | --- | --- |
| *Vigun05g248600* |  | 5’-CACTTGGCTACAATCCCTATCA -3’ | 5’-ACGGTCCAACCCAGTAAATC-3’ |
| *Vigun05g282700* |  | 5’-CACTGGTCTCTTCTACTGGATTG-3’ | 5’-GCAACACCGTGGATAGGAATA-3’ |
| *Vigun09g276100* | *VuTPS6* | 5’-CCGTCATCGGTCACAAGAAA-3’ | 5’-GTGCAGTAGACGAGAACGAAG-3’ |
| *Vigun06g141300* |  | 5’-CCATTGTTCGTGAAGTCGTGAG-3’ | 5’-GTGAGTTGTTGTTCGGCAGAG-3’ |
| *Vigun10g181500* |  | 5’-TCTCTGGTGGGTCGATGAA-3’ | 5’-CAATAAGAGGTCCAACCCAGTAG-3’ |
| *Vigun04g203000* | *VuACTIN* | 5’-TCAGGTGTCCAGAGGTGTTGTA-3’ | 5’-ATGGTTGTGCCTCCTGAAAGTA-3’ |
| *Vigun03g378000* | *VuTPS11* | 5’-AAGTATTGCTCGTTCGGTGTCTA-3’ | 5’-TGGATGAGGATGGTGTTGATGG-3’ |
| *Vigun03g186900* | *VuCYP707A-like* | 5’-ACTAGGTTGTCCATGCGTGAT-3’ | 5’-CTGATGCGACTGTGGTATTCTC-3’ |
| *Glyma.04g003200* |  | 5’-CATCACTGGTACCGGGATTAAC-3’ | 5’-CGAATGGTCCAACCCAGAATA-3’ |
| *Glyma.04g083200* |  | 5’-GCCGCAACAGCTTCTTATCT-3’ | 5’-CCCAAACTACTCCTTGACCATAC-3’ |
| *Glyma.08g015300* |  | 5’-ACTGTCATGGGTGTCAACAG-3’ | 5’-GCAGTAGACAAGGGCAAAGA-3’ |
| *Glyma.13g224900* |  | 5’-TCCCTTCCGGTTCTGATTTG-3’ | 5’-GCTCACCAATCGCTCTGTTA-3’ |
| *Glyma.19g181300* |  | 5’-CTAGGGCCTGCTGTCATATTC-3’ | 5’-CACGGACTGGTGGTAGAATG-3’ |
| *Glyma.02g091900* | *GmACTIN* | 5’-TCAGCCACACTGTCCCTATC-3’ | 5’-GCTCGTAGTCAAGGGCAATG-3’ |
