## Supplemental Data1 for "Understanding water conservation vs. profligation traits in vegetable legumes through a physio-transcriptomic-functional approach": Supplemental Data S1.R

```
###GLM###


if (!requireNamespace("BiocManager", quietly = TRUE))
  install.packages("BiocManager")

#BiocManager::install("edgeR")
#install("statmod")


#soybean
s_count2 <- read.table("soybean.genes.read.txt",header=T)
rownames(s_count2) <- s_count2[,1]
s_count1 <- s_count2[,-1]
#del_s <- which(rowSums(s_count1)<50)
#s_count <- s_count1[-del_s,]

#cowpea
c_count2 <- read.table("Cowpea.genes.read.txt",header=T)
rownames(c_count2) <- c_count2[,1]
c_count1 <- c_count2[,-1]
#del_c <- which(rowSums(c_count1)<50)
#c_count <- c_count1[-del_c,]


#Well watered=1,Mild drought=2,Severe drought=3,Recovery=4

treatment_s <- c(rep(2,9),rep(3,9),rep(1,9),rep(4,9))
treatment_c <- c(rep(2,9),rep(1,9),rep(3,9),rep(4,9))
tivec <- rep(c(12,12,12,16,16,16,6,6,6),4)

library(edgeR)
library(statmod)

#DE for soybean
# design matrix
design_s <- model.matrix(~treatment_s+tivec+treatment_s*tivec)
rownames(design_s) <- colnames(s_count1)

#Filter genes by expression level
keep_s <- filterByExpr(s_count1, design_s)
s_count <- s_count1[keep_s,]

#Normalize
d_s <- DGEList(s_count)
d_s <- calcNormFactors(d_s)

# dispersion coefficients

d_s <- estimateDisp(d_s, design_s, robust=TRUE)

#Fit the NB GLMs
fit_s <- glmFit(d_s, design_s)


# Likelihood ratio tests for each variable
#treatment_s
results_treat_s <- glmLRT(fit_s, coef=2)
treat_s_de <- topTags(results_treat_s,n=dim(s_count)[1])
write.csv(treat_s_de$table,file="soybean_de_treatment_all.csv")
treat_s_sig <- which(treat_s_de$table$FDR<0.05)
write.csv(treat_s_de$table[treat_s_sig,],file="soybean_de_treatment_fdr_005.csv")
treat_s1 <- treat_s_de$table[treat_s_sig,]
write.csv(treat_s1[which(abs(treat_s1[,1])>=2),],file="soybean_de_treatment_fdr_005_logfc_2.csv")

#time_s
results_time_s <- glmLRT(fit_s, coef=3)
time_s_de <- topTags(results_time_s,n=dim(s_count)[1])
write.csv(time_s_de$table,file="soybean_de_time_all.csv")
time_s_sig <- which(time_s_de$table$FDR<0.05)
write.csv(time_s_de$table[time_s_sig,],file="soybean_de_time_fdr_005.csv")
time_s1 <- time_s_de$table[time_s_sig,]
write.csv(treat_s1[which(abs(time_s1[,1])>=2),],file="soybean_de_time_fdr_005_logfc_2.csv")

#interaction effect for soybean
results_inter_s <- glmLRT(fit_s, coef=4)
inter_s_de <- topTags(results_inter_s,n=dim(s_count)[1])
write.csv(inter_s_de$table,file="soybean_de_inter_all.csv")
inter_s_sig <- which(inter_s_de$table$FDR<0.05)
write.csv(inter_s_de$table[inter_s_sig,],file="soybean_de_inter_fdr_005.csv")
inter_s1 <- inter_s_de$table[inter_s_sig,]
write.csv(inter_s1[which(abs(inter_s1[,1])>=2),],file="soybean_de_inter_fdr_005_logfc_2.csv")


#DE for cowpea
# design matrix
design_c <- model.matrix(~treatment_c+tivec+treatment_c*tivec)
rownames(design_c) <- colnames(c_count1)

#Filter genes by expression level
keep_c <- filterByExpr(c_count1, design_c)
c_count <- c_count1

#Normalize
d_c <- DGEList(c_count)
d_c <- calcNormFactors(d_c)

# dispersion coefficients

#d_c <- estimateDisp(d_c, design_c, robust=TRUE)

#Fit the NB GLMs
fit_c <- glmFit(d_c, design_c,dispersion=0.1)


# Likelihood ratio tests for each variable
#treatment_c
results_treat_c <- glmLRT(fit_c, coef=2)
treat_c_de <- topTags(results_treat_c,n=dim(c_count)[1])
write.csv(treat_c_de$table,file="cowpea_de_treatment_all.csv")
treat_c_sig <- which(treat_c_de$table$FDR<0.05)
write.csv(treat_c_de$table[treat_c_sig,],file="cowpea_de_treatment_fdr_005.csv")
treat_c1 <- treat_c_de$table[treat_c_sig,]
write.csv(treat_c1[which(abs(treat_c1[,1])>=2),],file="cowpea_de_treatment_fdr_005_logfc_2.csv")

#time_c
results_time_c <- glmLRT(fit_c, coef=3)
time_c_de <- topTags(results_time_c,n=dim(c_count)[1])
write.csv(time_c_de$table,file="cowpea_de_time_all.csv")
time_c_sig <- which(time_c_de$table$FDR<0.05)
write.csv(time_c_de$table[time_c_sig,],file="cowpea_de_time_fdr_005.csv")
time_c1 <- time_c_de$table[time_c_sig,]
write.csv(treat_c1[which(abs(time_c1[,1])>=2),],file="cowpea_de_time_fdr_005_logfc_2.csv")

#interaction effect for cowpea
results_inter_c <- glmLRT(fit_c, coef=4)
inter_c_de <- topTags(results_inter_c,n=dim(c_count)[1])
write.csv(inter_c_de$table,file="cowpea_de_inter_all.csv")
inter_c_sig <- which(inter_c_de$table$FDR<0.05)
write.csv(inter_c_de$table[inter_c_sig,],file="cowpea_de_inter_fdr_005.csv")
inter_c1 <- inter_c_de$table[inter_c_sig,]
write.csv(inter_c1[which(abs(inter_c1[,1])>=2),],file="cowpea_de_inter_fdr_005_logfc_2.csv")

####Cluster


#library(mclust) 
#library(cluster)
#library(NbClust)

Clust.kmeans = function(dat,standard=TRUE,nclust=NULL,
                        max.clust=50,rss.thresh=0.75,iter.max=1000,nstart=10,seeds=1234){
  if(!is.matrix(dat)){
    dat = as.matrix(dat)
  }
  # gene-wise standardization
  if(standard){
    dat = t(apply(dat,1,function(x) (x-mean(x,na.rm=TRUE))/sd(x,na.rm=TRUE)))
  }
  
  
  if(is.null(nclust)){
    for(nclust in 2:max.clust){
      set.seed(seeds)
      clust.res = kmeans(dat,centers=nclust,iter.max=iter.max,nstart=nstart)
      rss.ratio = 1-clust.res$tot.withinss/clust.res$totss
      if(rss.ratio >= rss.thresh){
        break
      }
    }
    if(nclust == max.clust & rss.ratio < rss.thresh){
      cat("Warning: the maximum number of clusters reached before the percentage of variance explained exceeds threshold! \n")
    }
  }else{
    set.seed(seeds)
    clust.res = kmeans(dat,centers=nclust,iter.max=iter.max,nstart=nstart)
  }
  
  list(nclust=nclust,clust=clust.res$cluster,center=clust.res$centers,totss=clust.res$totss,
       withinss=clust.res$withinss,tot.withinss=clust.res$tot.withinss,betweenss=clust.res$betweenss,
       size=clust.res$size,dat.clust=dat)
}


cluster_plot <- function(ex,cu,ci=1:12,w=12,h=16,file="s_treat_de_cu.pdf"){
  
  GD <- log10(ex+1)
  nti <- c(6,12,16,18,24,28,30,36,40,42,48,52)
  pdf(file,height=16,width=12)
  par(mar=c(1,1,2,2),oma=c(5,5,1,5),mfrow=c(6,2))
  for(i in ci){
    index <- which(cu==i)
    cu_GD <- GD[index,]
    plot(0,0,pch=16,type="n",col="#757575",xlab=" ",ylab=" ",xlim=c(5,53),ylim=c(-0.05,round(max(cu_GD))+0.1),xaxt="n",yaxt="n",xaxs="i", yaxs="i")
    for(ii in 1:length(index)){
      lines(nti[1:3],cu_GD[ii,1:3],lwd=1,col="#C4C4C450")
      lines(nti[4:6],cu_GD[ii,4:6],lwd=1,col="#C4C4C450")
      lines(nti[7:9],cu_GD[ii,7:9],lwd=1,col="#C4C4C450")
      lines(nti[10:12],cu_GD[ii,10:12],lwd=1,col="#C4C4C450")
    }
    lines(nti[1:3],colMeans(GD[index,1:3]),lwd=3,col="#fa4685")
    lines(nti[4:6],colMeans(GD[index,4:6]),lwd=3,col="#fa4685")
    lines(nti[7:9],colMeans(GD[index,7:9]),lwd=3,col="#fa4685")
    lines(nti[10:12],colMeans(GD[index,10:12]),lwd=3,col="#fa4685")
    segments(17,-10,17,20,lwd=2,col="#436EEE")
    segments(29,-10,29,20,lwd=2,col="#436EEE")
    segments(41,-10,41,20,lwd=2,col="#436EEE")
    if(i>=(max(ci)-1)){
      axis(1,nti,rep(c(6,12,16),4),mgp=c(2.7,1.6,0),cex.axis=1.5,las=1,tck=-.04)
      axis(1,nti[c(4,7,10)],rep(c(6,12,16),4)[c(4,7,10)],mgp=c(2.7,1.6,0),cex.axis=1.5,las=1,tck=-.04)
    }
    mtext(paste("Cluster",i,sep=""),3,cex=1.5,line=0.50)
    text(c(11,23,35,47),(round(max(cu_GD))+0.1)*0.95,c("WW","MD","SD","R"),font=1,cex=1.5)
    if(i==(max(ci)-1)){
      mtext("Time (Day)",1,cex=1.5,line=3.50,adj=1.2)
    }
    if(i%%2==1){
      axis(2,seq(0,round(max(cu_GD))+0.1,0.5),seq(0,round(max(cu_GD))+0.1,0.5),
           mgp=c(2.7,1.6,0),cex.axis=1.5,las=1,tck=-.04)
    }
    if(i%%2==0){
      axis(4,seq(0,round(max(cu_GD))+0.1,0.5),seq(0,round(max(cu_GD))+0.1,0.5),
           mgp=c(2.7,1.6,0),cex.axis=1.5,las=1,tck=-.04)
    }
    if(i==ci[5]){
      mtext(expression('log'[10]*'(FPKM+1)'),2,cex=1.5,line=3.50,adj=-2)
    }
    if(i==ci[6]){
      mtext(expression('log'[10]*'(FPKM+1)'),4,cex=1.5,line=5,adj=-2)
    }
  }
  
  
  dev.off()
  
  
}


#FPKM
s3 <- read.csv("Soybean-ZN6-genes.fpkm.anno.csv",header = F,fill=T)
s31 <- s3[,1]
s32 <- s3[,-1]
s2 <- s32[-c(1:2),c(7:9,1:6,16:18,10:15,25:27,19:24,34:36,28:33)]
s1 <- apply(s2,2,as.numeric)
rownames(s1) <- s31[-c(1,2)]

c3 <- read.csv("Cowpea-TZ30-genes.fpkm.anno.csv",header = F,fill=T)
c31 <- c3[,1]
c32 <- c3[,-1]
c2 <- c32[-c(1,2),c(7:9,1:6,16:18,10:15,25:27,19:24,34:36,28:33)]
c1 <- apply(c2,2,as.numeric)
rownames(c1) <- c31[-c(1,2)]

#de

s_treat11 <- read.csv("./soybean/soybean_de_treatment_fdr_005.csv")
s_treat <- s_treat11[which(abs(s_treat11$logFC)>1),]

s_time11 <- read.csv("./soybean/soybean_de_time_fdr_005.csv")
s_time <- s_time11[which(abs(s_time11$logFC)>0.5),]

s_inter11 <- read.csv("./soybean/soybean_de_inter_fdr_005.csv")
s_inter <- s_inter11[which(abs(s_inter11$logFC)>0.25),]

c_treat11 <- read.csv("./cowpea/cowpea_de_treatment_fdr_005.csv")
c_treat <- c_treat11[which(abs(c_treat11$logFC)>1),]

c_time11 <- read.csv("./cowpea/cowpea_de_time_fdr_005.csv")
c_time <- c_time11[which(abs(c_time11$logFC)>0.5),]

c_inter11 <- read.csv("./cowpea/cowpea_de_inter_fdr_005.csv")
c_inter <- c_inter11[which(abs(c_inter11$logFC)>0.25),]
#DE_ex

s_treat_ex <- s1[match(s_treat[,1],rownames(s1)),]
s_time_ex <- s1[match(s_time[,1],rownames(s1)),]
s_inter_ex <- s1[match(s_inter[,1],rownames(s1)),]

c_treat_ex <- c1[match(c_treat[,1],rownames(c1)),]
c_time_ex <- c1[match(c_time[,1],rownames(c1)),]
c_inter_ex <- c1[match(c_inter[,1],rownames(c1)),]

s_treat_em <- c();s_time_em <- c();s_inter_em <- c();
c_treat_em <- c();c_time_em <- c();c_inter_em <- c();
for(i in 1:12){
  s_treat_em <- cbind(s_treat_em,rowMeans(s_treat_ex[,(3*i-2):(3*i)]))
  s_time_em <- cbind(s_time_em,rowMeans(s_time_ex[,(3*i-2):(3*i)]))
  s_inter_em <- cbind(s_inter_em,rowMeans(s_inter_ex[,(3*i-2):(3*i)]))
  c_treat_em <- cbind(c_treat_em,rowMeans(c_treat_ex[,(3*i-2):(3*i)]))
  c_time_em <- cbind(c_time_em,rowMeans(c_time_ex[,(3*i-2):(3*i)]))
  c_inter_em <- cbind(c_inter_em,rowMeans(c_inter_ex[,(3*i-2):(3*i)]))
}

###number of cluster####

#s_treat=12
s_treat_cu <- Clust.kmeans(dat=s_treat_em,max.clust=130,rss.thresh=0.75)
write.csv(cbind(s_treat,cluster=s_treat_cu$clust),file="S_treat_DE_cluster.csv")
cluster_plot(ex=s_treat_em,cu=s_treat_cu$clust,ci=1:12,file="s_treat_de_cu_1_12.pdf")

#s_time=9
s_time_cu <- Clust.kmeans(dat=s_time_em,max.clust=130,rss.thresh=0.75)
write.csv(cbind(s_time,cluster=s_time_cu$clust),file="S_time_DE_cluster.csv")
cluster_plot_bug1(ex=s_time_em,cu=s_time_cu$clust,ci=1:9,file="s_time_de_cu.pdf")

#s_inter=7
s_inter_cu <- Clust.kmeans(dat=s_inter_em,max.clust=130,rss.thresh=0.75)
write.csv(cbind(s_inter,cluster=s_inter_cu$clust),file="S_inter_DE_cluster.csv")
cluster_plot_bug1(ex=s_inter_em,cu=s_inter_cu$clust,ci=1:7,file="s_inter_de_cu.pdf")

#c_treat=21
c_treat_cu <- Clust.kmeans(dat=c_treat_em,max.clust=130,rss.thresh=0.73)
write.csv(cbind(c_treat,cluster=c_treat_cu$clust),file="C_treat_DE_cluster.csv")
cluster_plot(ex=c_treat_em,cu=c_treat_cu$clust,ci=1:12,file="c_treat_de_cu_1_12.pdf")
cluster_plot_bug1(ex=c_treat_em,cu=c_treat_cu$clust,ci=13:21,file="c_treat_de_cu_13_21.pdf")

#c_time=12
c_time_cu <- Clust.kmeans(dat=c_time_em,max.clust=130,rss.thresh=0.75)
write.csv(cbind(c_time,cluster=c_time_cu$clust),file="C_time_DE_cluster.csv")
cluster_plot(ex=c_time_em,cu=c_time_cu$clust,ci=1:12,file="c_time_de_cu.pdf")


#c_inter=39
c_inter_cu <- Clust.kmeans(dat=c_inter_em,max.clust=130,rss.thresh=0.75)
write.csv(cbind(c_inter,cluster=c_inter_cu$clust),file="C_inter_DE_cluster.csv")
cluster_plot_bug1(ex=c_inter_em,cu=c_inter_cu$clust,ci=1:6,file="c_inter_de_cu.pdf")

###inter-network


get_net_matrix <- function(em){
  
  library(glmnet)
  em1 <- t(em)
  nn <- dim(em1)[2]
  netm <- matrix(0,nn,nn)
  for(i in 1:nn){
    xs <- em1[,-i]
    ys <- em1[,i]
    cv <- cv.glmnet(x = xs, y = ys,alpha = 0)
    best_coef <- as.numeric(coef(cv, s = cv$lambda.min))[-1]
    fit_res <- cv.glmnet(x = xs, y = ys,alpha =1,
                         penalty.factor = 1/best_coef,
                         keep = TRUE)
    best_alasso_coef1 <- coef(fit_res, s = fit_res$lambda.min)
    netm[i,-i] <- as.numeric(best_alasso_coef1[-1])
  }
  return(netm)
}

s3 <- read.csv("../../Soybean-ZN6-genes.fpkm.anno.csv",header = F,fill=T)
s31 <- s3[,1]
s32 <- s3[,-1]
s2 <- s32[-c(1:2),c(7:9,1:6,16:18,10:15,25:27,19:24,34:36,28:33)]
s1 <- apply(s2,2,as.numeric)
rownames(s1) <- s31[-c(1,2)]

c3 <- read.csv("../../Cowpea-TZ30-genes.fpkm.anno.csv",header = F,fill=T)
c31 <- c3[,1]
c32 <- c3[,-1]
c2 <- c32[-c(1,2),c(7:9,1:6,16:18,10:15,25:27,19:24,34:36,28:33)]
c1 <- apply(c2,2,as.numeric)
rownames(c1) <- c31[-c(1,2)]


ms2 <- read.csv("s_gene_module.csv")
mc2 <- read.csv("c_gene_module.csv")

sc <- names(table(ms2[,2]))
cc <- names(table(mc2[,2]))


msem <- s1[match(ms2[,1],rownames(s1)),]
mcem <- c1[match(mc2[,1],rownames(c1)),]

scm <- c()
for(i in 1:length(sc)){
  scm <- rbind(scm,colMeans(msem[which(ms2[,2]==sc[i]),]))
}


ccm <- c()
for(i in 1:length(cc)){
  ccm <- rbind(ccm,colMeans(mcem[which(mc2[,2]==cc[i]),]))
}

sml <- log10(scm+1)
cml <- log10(ccm+1)


s_net <- get_net_matrix(em=sml)
c_net <- get_net_matrix(em=cml)

library(igraph)


nework_igraph <- function(M,label,thred=0.1,lc="grey",file="module-network.pdf"){
  
  net <- M
  
  net[which(abs(net)<thred)] <- 0
  nn <- dim(net)[1]
  ne.mut <- sum(net!=0)
  index <- which(c(t(net))!=0)
  nge <- which(c(t(net))[index]<0)
  g1 <- graph.adjacency((net),mode="directed",weighted=T)
  allshape <- rep("circle",nn)
  allsize <- rep(17,nn)
  V(g1)$label <- paste("M",1:nn,sep="")
  e.w <<-abs(c(t(net))[index])*2
  nr <-  rep("triangle",length(e.w))
  t1 <- rep("#FF000050",length(nr))
  t1[nge] <- rep("#0000FF50",length(nge))
  coll <- label
  rr1 <<- nr
  #allcol[which(igraph::degree(g1)>10)] <- "palegreen3"
  pdf(file=file,height=8,width=8)
  set.seed(14)
  plot(g1,xlim=c(-1,1),ylim=c(-1,1),
       vertex.size=allsize,vertex.label.cex=1.2,edge.color=t1,
       edge.arrow.size=1,vertex.color=coll,edge.width=e.w,vertex.shape=allshape,
       vertex.frame.color="#B5B5B5",vertex.label.color=lc,layout=layout.lgl)
  dev.off()
  
}

nework_igraph(M=s_net,label=sc,thred=0.2,file="module-network_s.pdf",lc="yellow")
nework_igraph(M=c_net,label=cc,thred=0.2,file="module-network_c.pdf",lc="darkblue")


write.csv(cbind(color=sc,Label=paste("N",1:length(sc))),file="S_network.csv")
write.csv(cbind(color=cc,Label=paste("N",1:length(cc))),file="C_network.csv")
```
