## Supplemental Fig 1-12 for "Understanding water conservation vs. profligation traits in vegetable legumes through a physio-transcriptomic-functional approach"

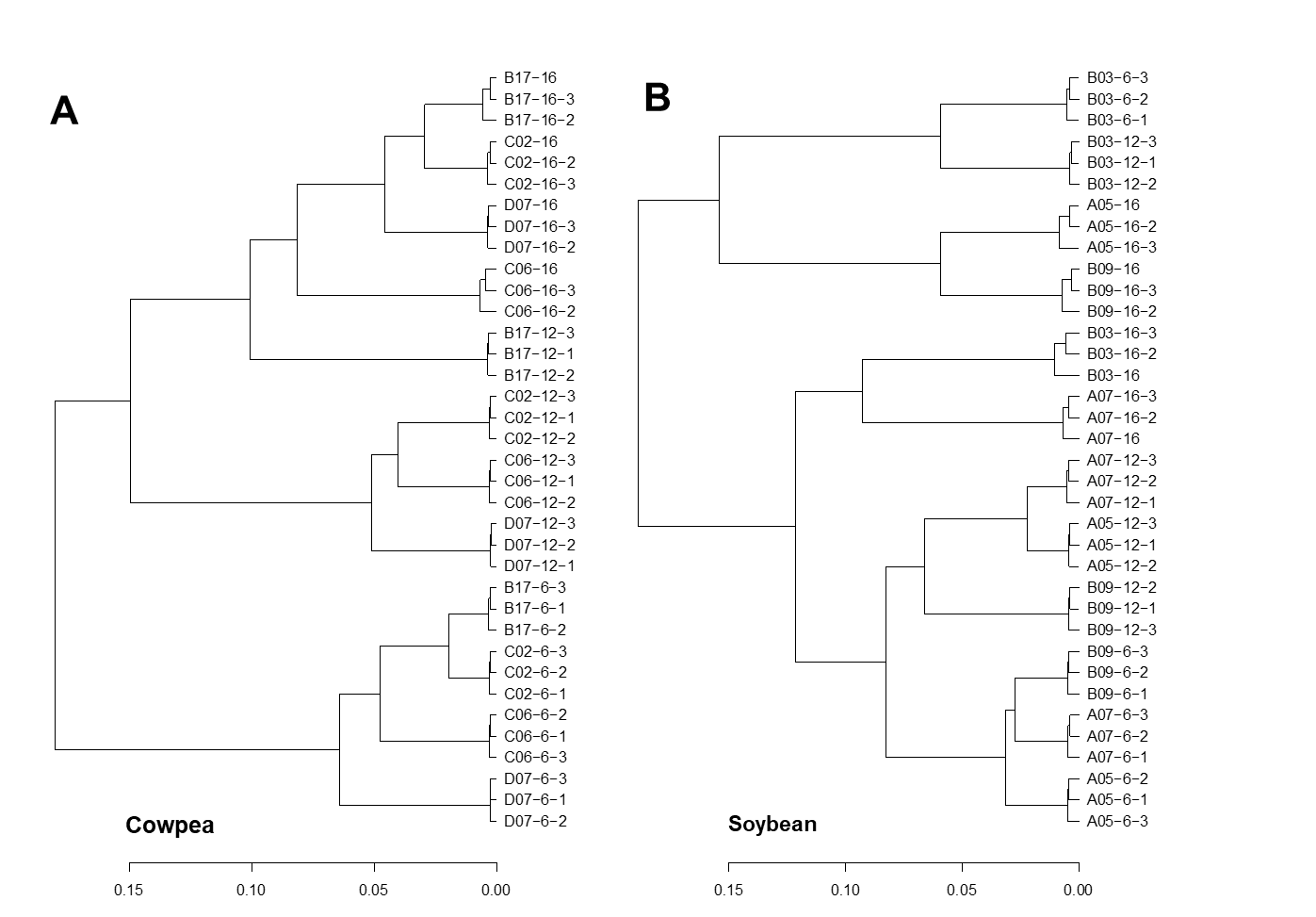

**A**

**B**

**Fig. S1.** Hierarchical clustering of the RNA-Seq data. A, C02, B17, C06 and D07 denote cowpea samples collected at the well-watered (WW), moderate soil drought (MD), severe soil drought (SD) and recovery (RC) stages, respectively; B, B03, A05, A07 and B09 denote soybean samples collected at WW, MD, SD and RC stages, respectively. The numbers -6, -12, and -16 denote time of day (6 am, 12 pm and 4 pm), and 1, 2, 3 denotes replicate number.

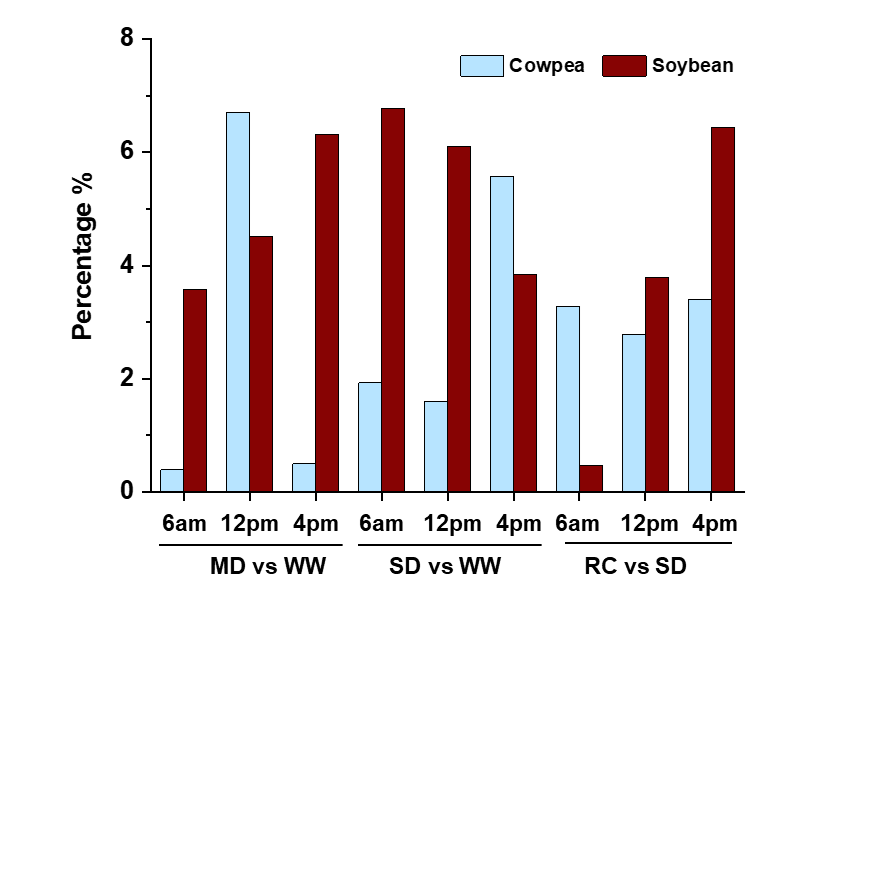

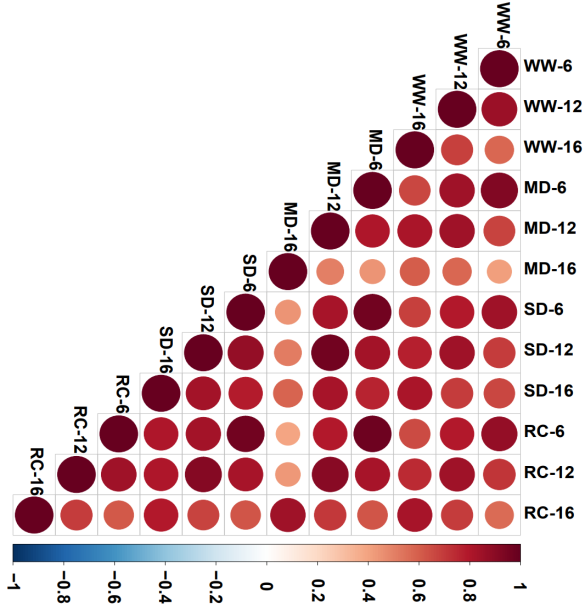
A
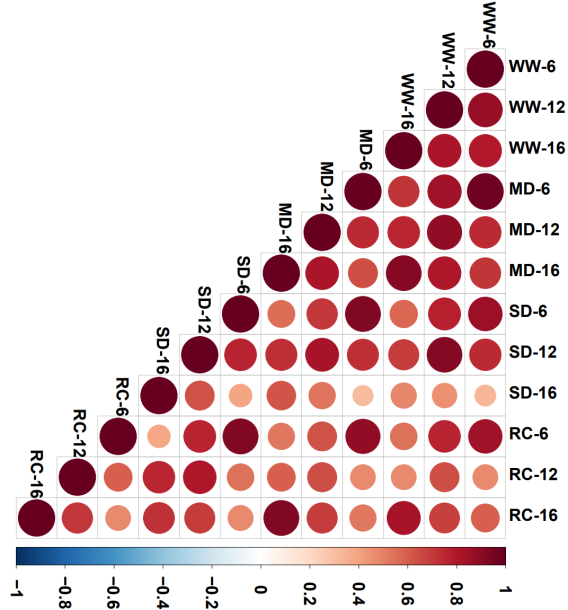

**Cowpea**

**Soybean**

**B**

**A**

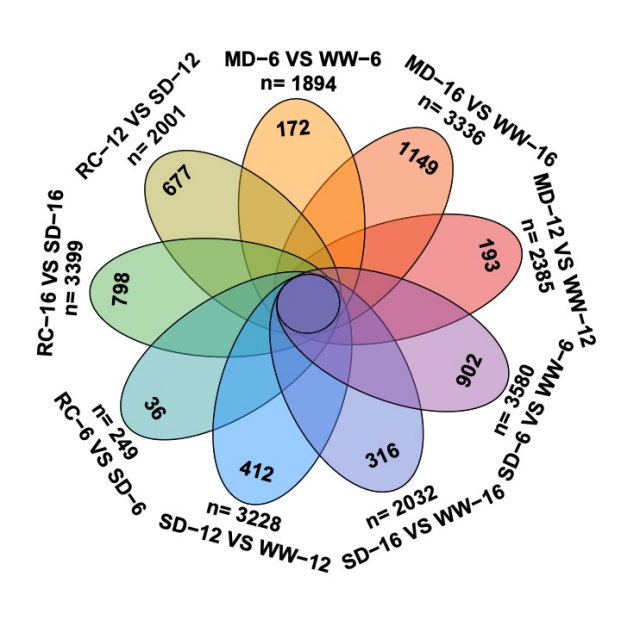

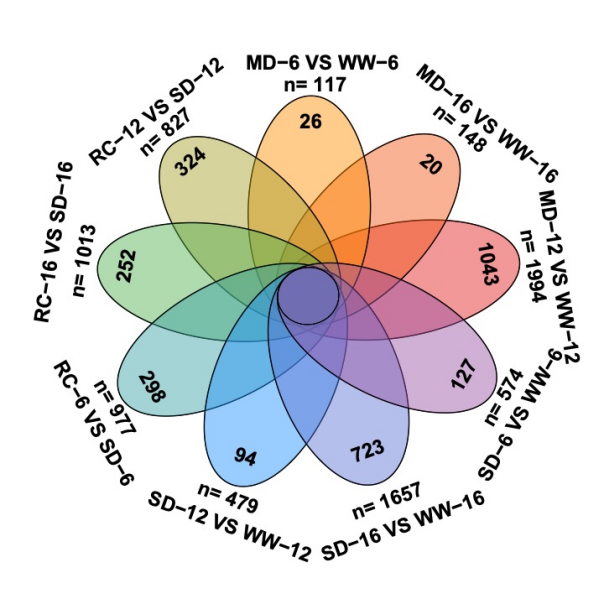

**C**

**Cowpea**

**Soybean**

**Fig. S2.** Overview of the transcriptomic data. A, Spearman correlation coefficiencies between samples. B, Percentage of DEGs relative to the total gene number of cowpea or soybean in each comparison. C, Number of DEGs in each comparison. The total and comparison-specific numbers were shown. WW: well-watered control; MD: moderate soil drought; SD: severe soil drought. RC: recovery phase. The numbers -6, -12, and -16 denote time of day (6 am, 12 pm and 4 pm).

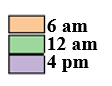

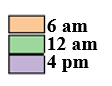

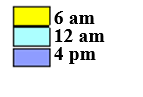

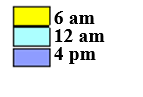

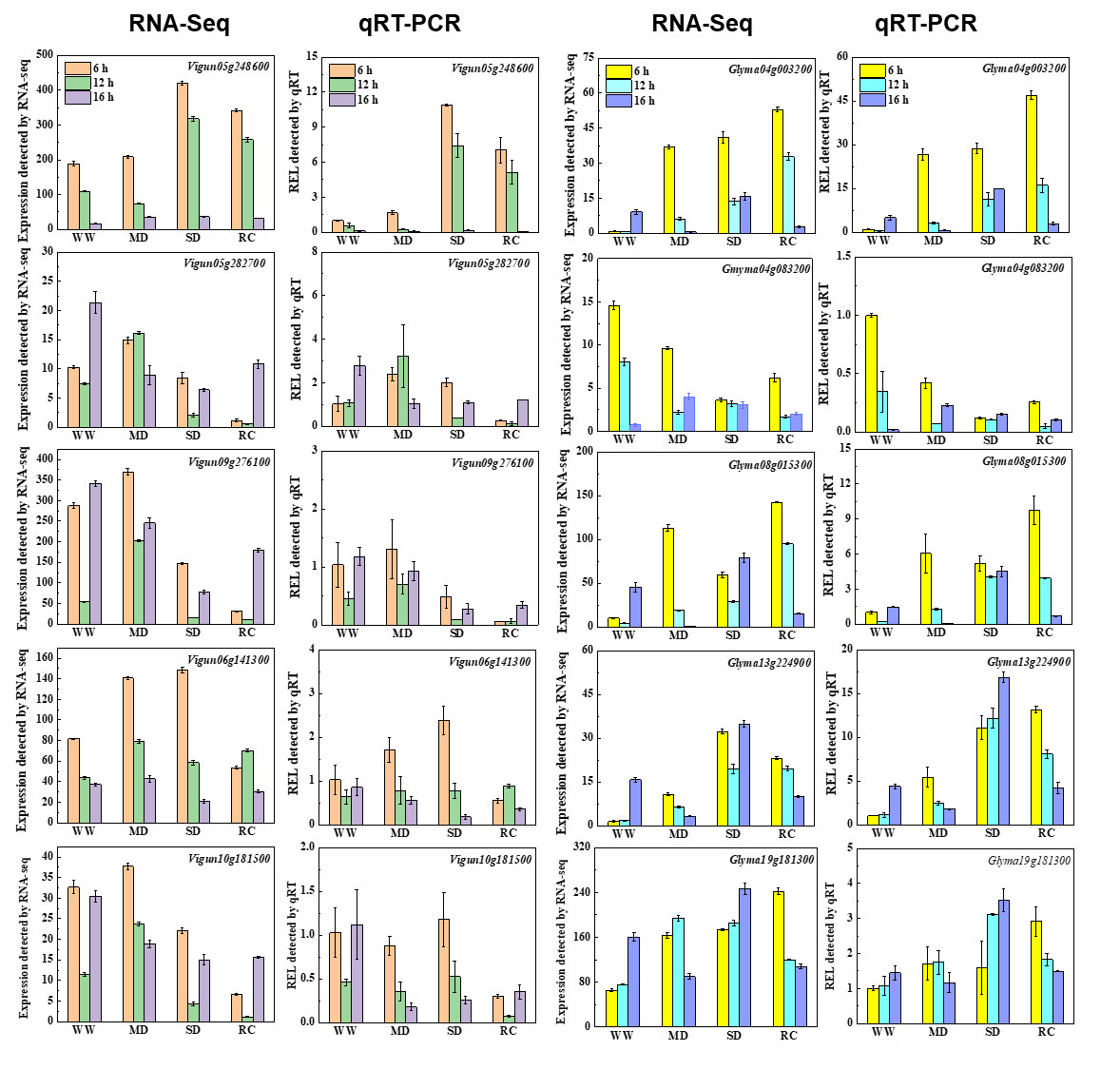

**Soybean**

**Cowpea**

**Fig. S3.** Validation of RNA-Seq by qRT-PCR on 10 randomly selected genes from the two crops. Data are the means of three replicates (±SD). WW: well-watered control; MD: moderate soil drought; SD: severe soil drought. RC: recovery phase.

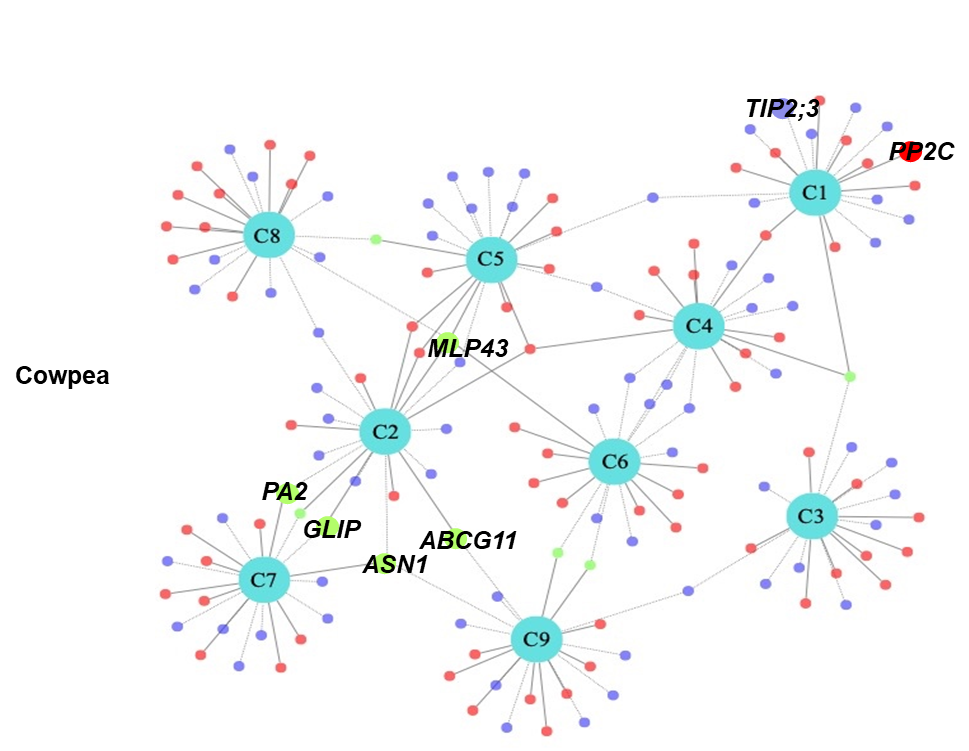

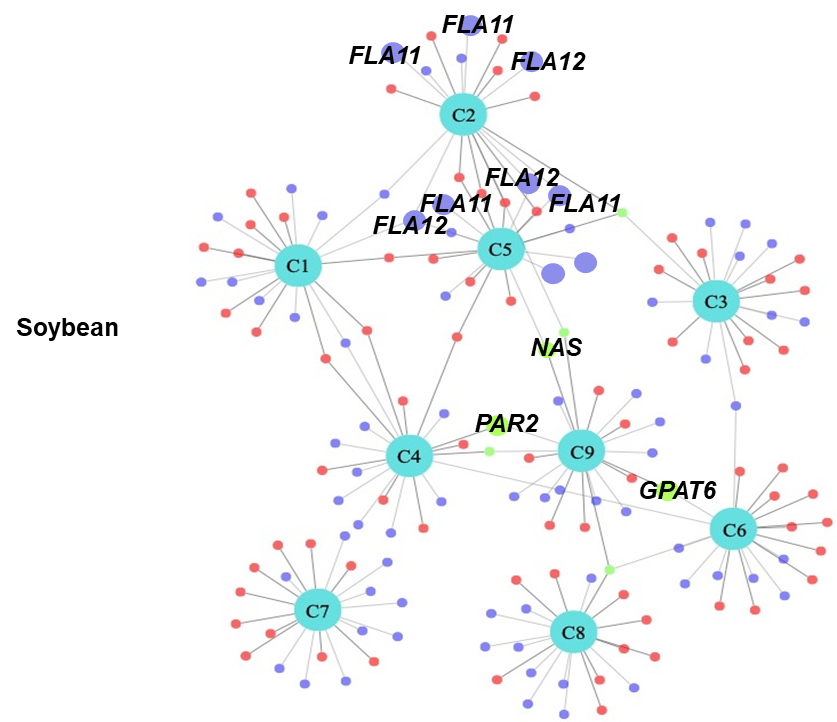

**Fig. S4.** Top up- and down-regulated genes in the nine comparisons and their topological relationships. The networks showing the topological relationship of the top DEGs in various pairwise comparisons were generated using Cytoskype based on the pairwise relationship between them. C1-C9: nine comparisons. C1: MD-6 am vs WW-6 am; C2: MD-12 pm vs WW-12 pm; C3: MD-4 pm vs WW-4 pm; C4: SD-6 am vs WW-6 am; C5: SD-12 pm vs WW-12 pm; C6 : SD-4 pm vs WW-4 pm; C7: RC-6 am vs SD-6 am; C8: RC-12 pm vs SD-12 pm; C9: RC-4 pm vs SD-4 pm. Red dots: up-regulated genes; blue dots: down-regulated genes; green dot: genes that were down-regulated in one comparison but up-regulated in another. WW: well-watered control; MD: moderate soil drought; SD: severe soil drought. RC: recovery phase.

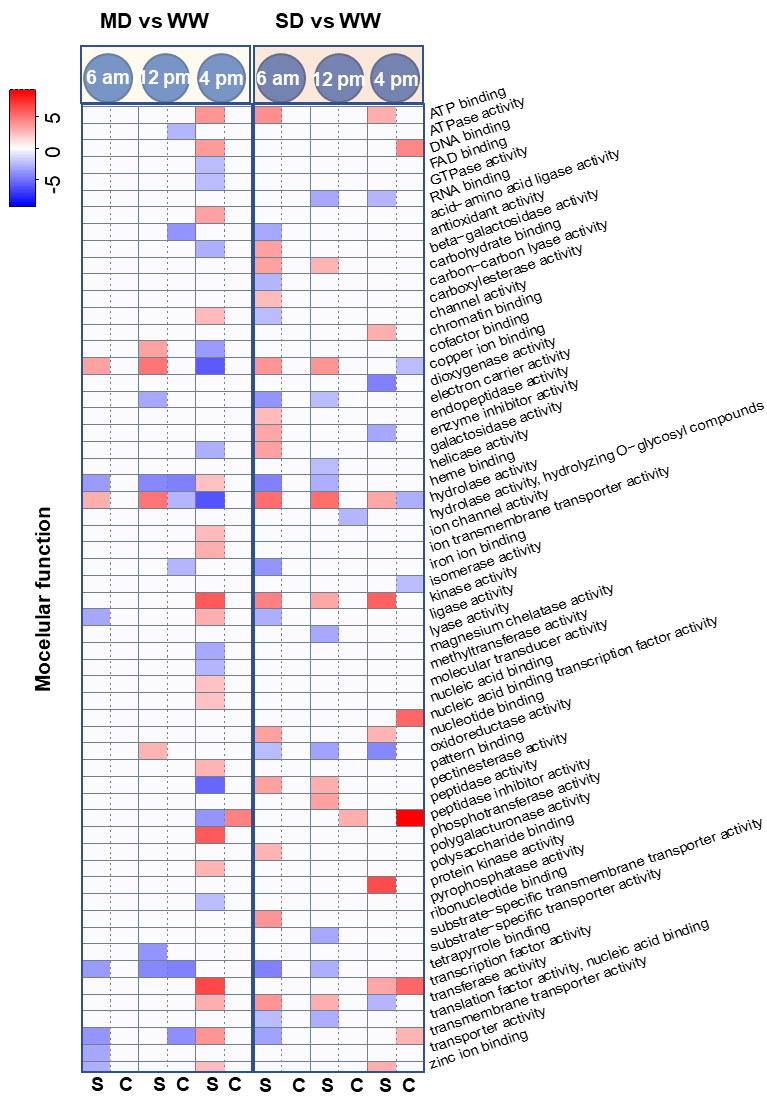
**Fig. S5A.** Gene ontology enrichment analysis on molecular function of the DEGs under moderate and severe soil drought conditions in comparison to the well-watered condition. WW: well-watered control; MD: moderate soil drought; SD: severe soil drought; S: soybean; C: cowpea.

**Z-score**

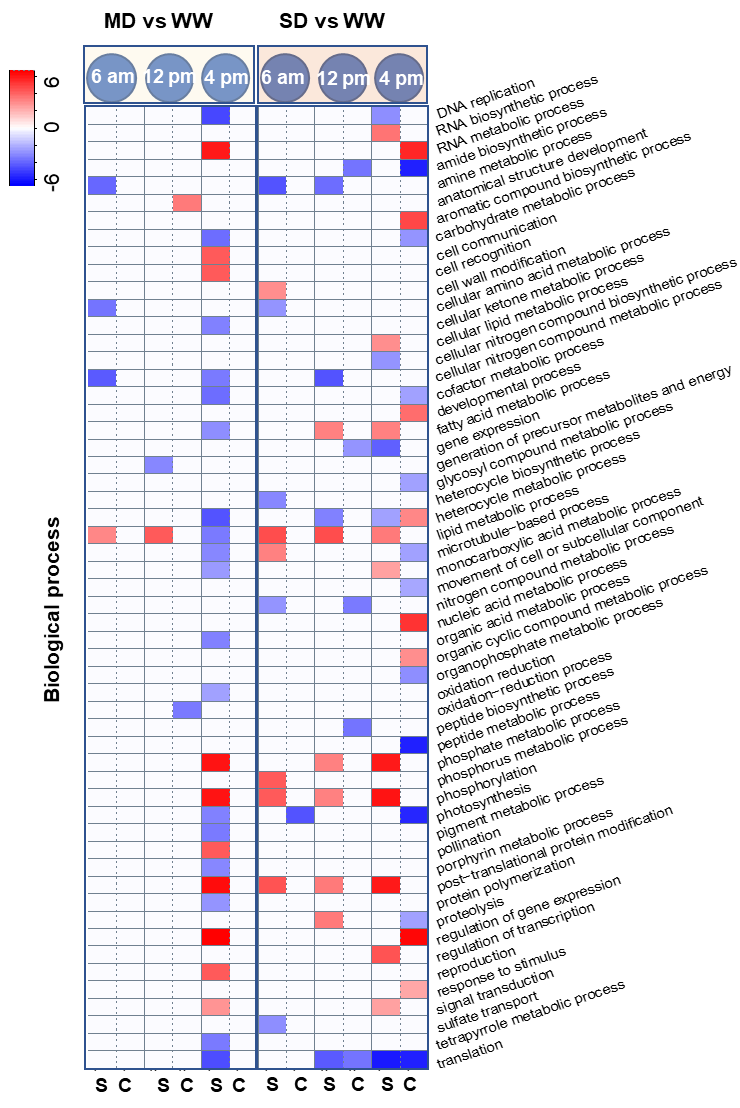

**Z-score**

**Fig. S5B.** Gene ontology enrichment analysis on biological process of the DEGs under moderate and severe soil drought conditions in comparison to the well-watered condition. WW: well-watered control; MD: moderate soil drought; SD: severe soil drought; S: soybean; C: cowpea.

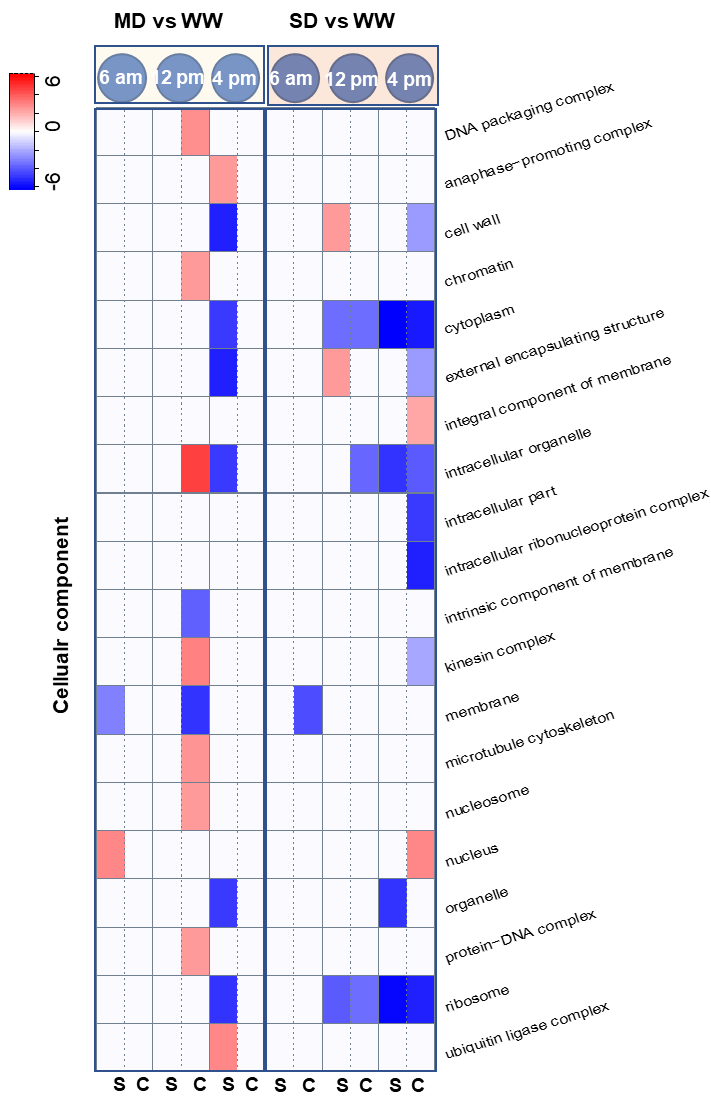

**Z-score**

**Fig. S5C.** Gene ontology enrichment analysis on cellular component of the DEGs under moderate and severe soil drought conditions in comparison to the well-watered condition. WW: well-watered control; MD: moderate soil drought; SD: severe soil drought; S: soybean; C: cowpea.

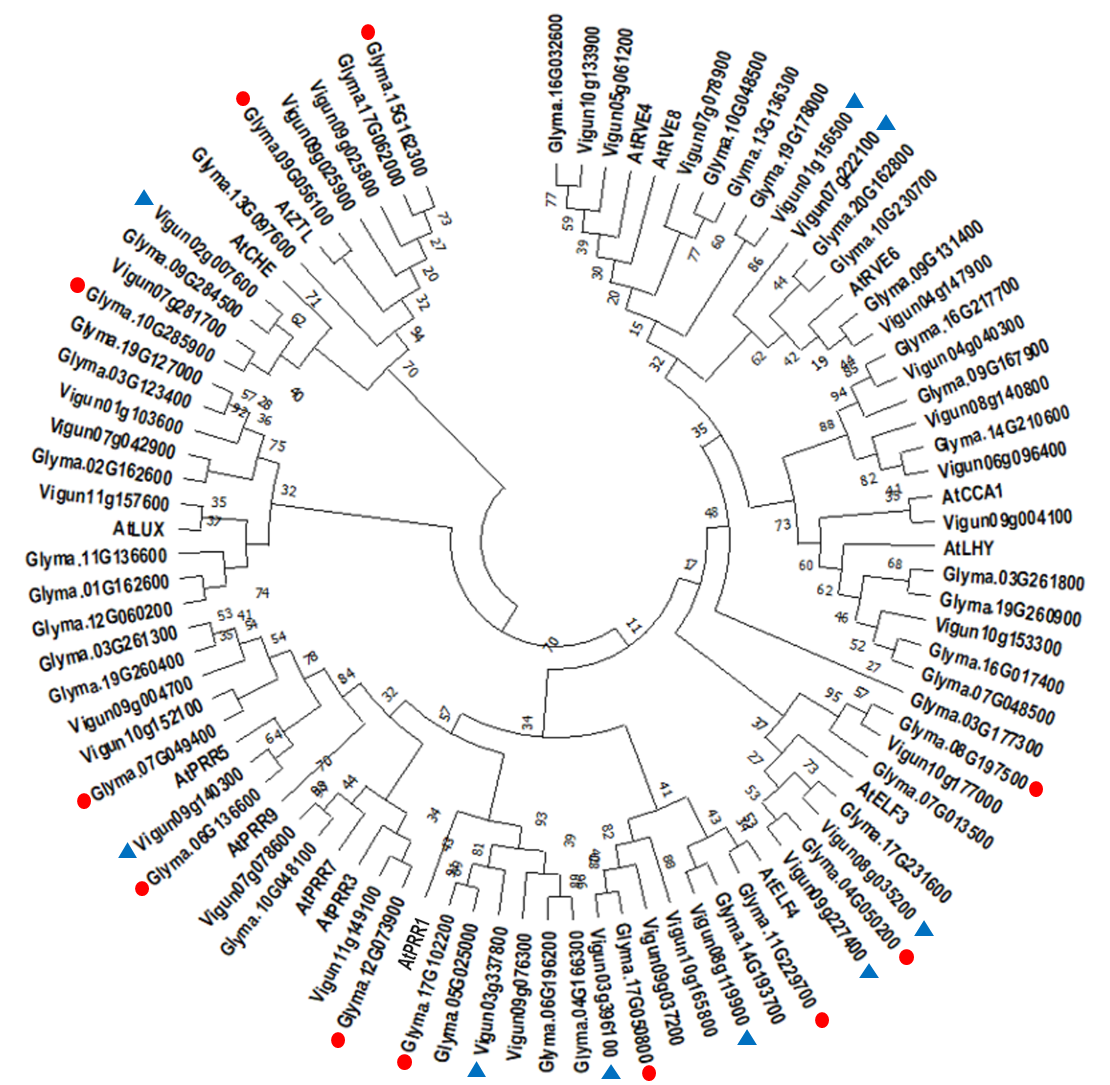

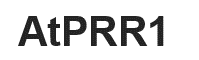

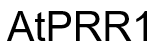

**Fig. S6.** Phylogenetic tree of the orthologs of core circadian clock genes from *Arabidopsis thaliana*, *Glycine max* and *Vigna unguiculata*. The phylogeny was constructed by using the maximum-likelihood method with 1000 bootstrap replicates. Genes showing changes of amplitude or trend in expression under progressive soil drought were marked by red circle (*Glycine max*) and blue triangle (*Vigna unguiculata*), respectively.

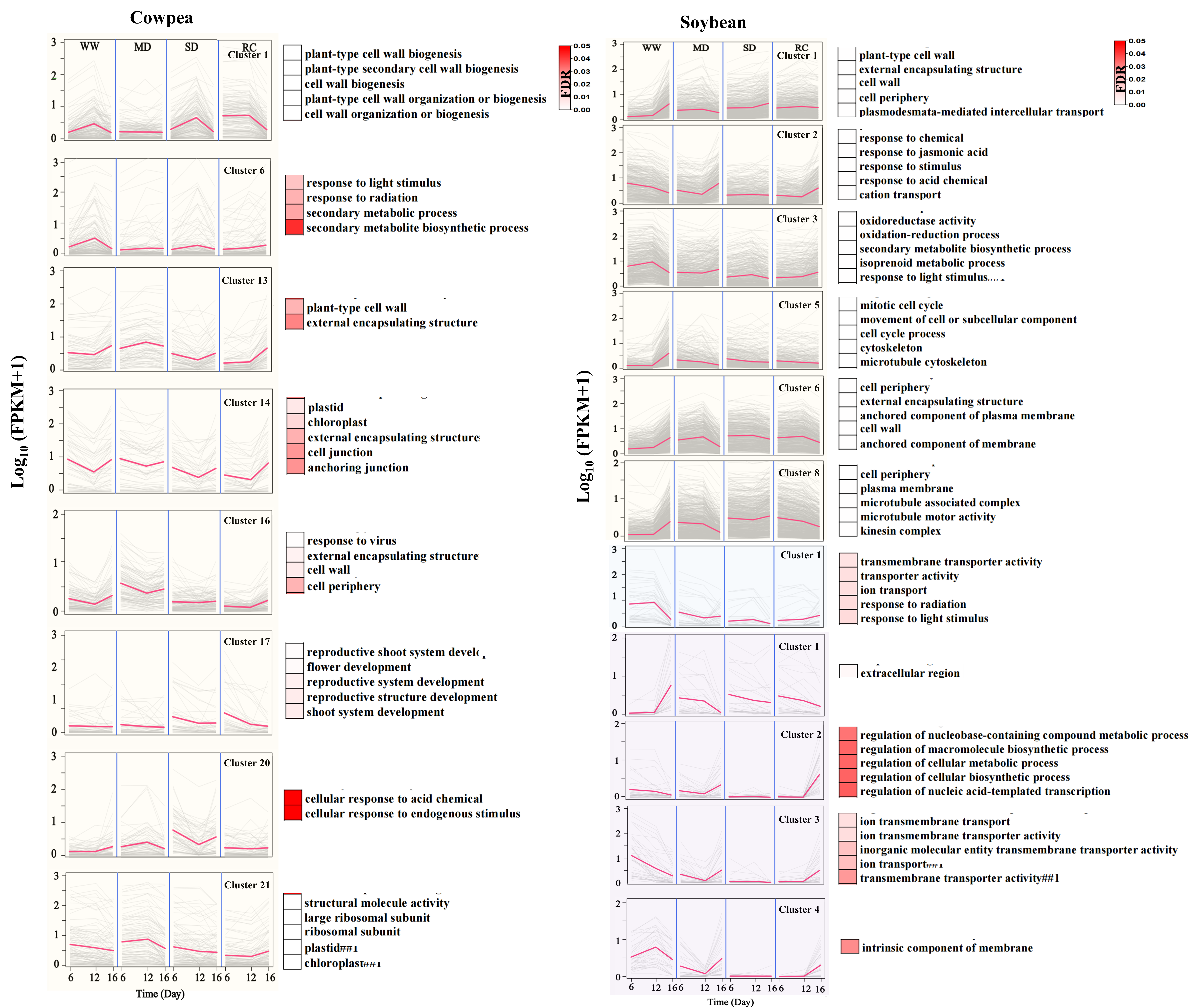

**Fig. S7.** Gene clusters and their functional annotations in the two crops. Differentially expressed genes identified from the generalized linear model (GLM)-based analysis were clustered using the k-means method. The background color of the cluster, namely light yellow, light blue or light purple, indicates that the cluster was affected by drought, TOD or their interactions, respectively. The top five or all (if less than five) enriched GO terms based on false discovery rate (FDR) were shown for each cluster. Only those with an FDR ≤ 0.05 were considered. WW: well-watered control; MD: moderate soil drought; SD: severe soil drought, RC: recovery phase.

**Dynamic Tree Cut**

**Height**

**Merged dynamic**

**Merged dynamic**

**Cluster Dendrogram**

**Height**

**Dynamic Tree Cut**

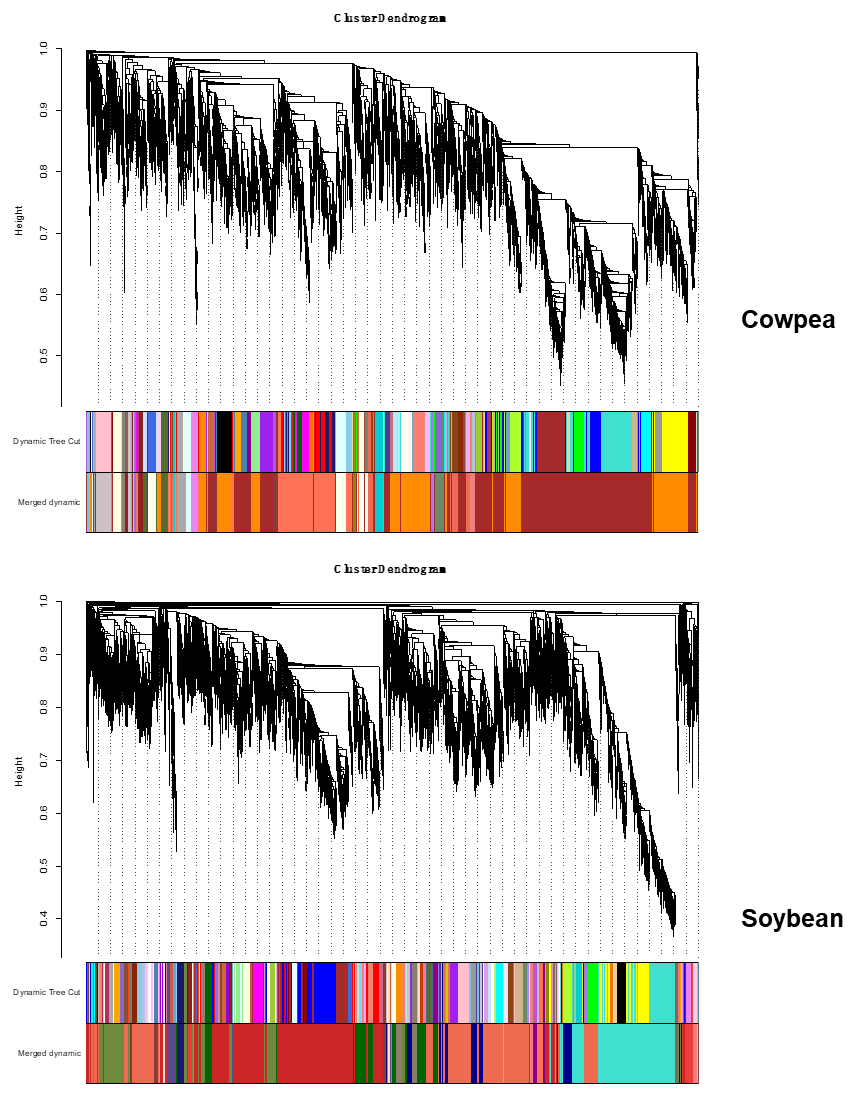

**Cluster Dendrogram**

**Fig. S8.** Hierarchical cluster trees showing the modules of coexpressed genes. Each of the modules is represented by a major tree branch.

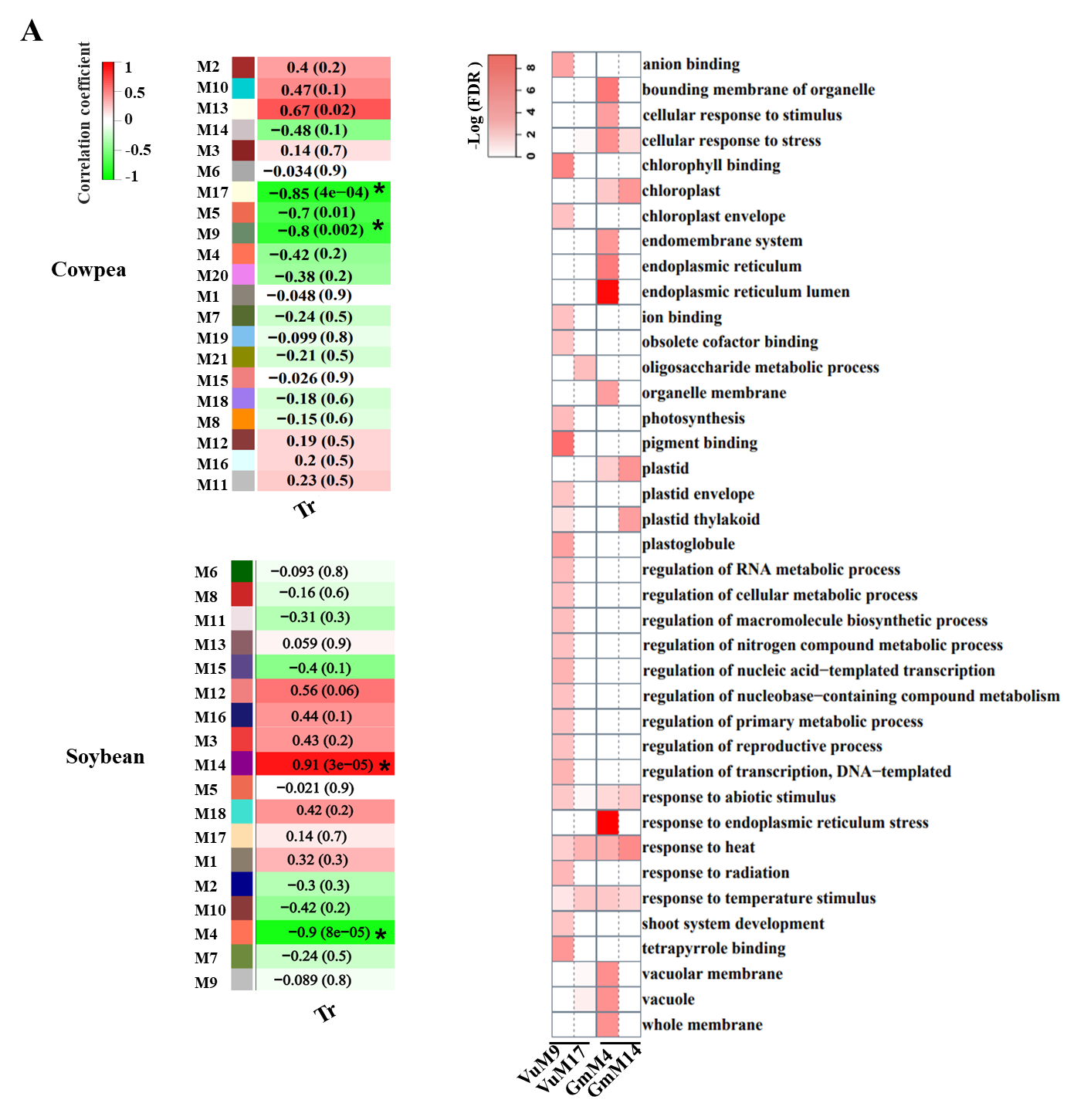

**B**

**Fig. S9.** Modules associated with transpiration rate in cowpea and soybean. A, The trait association module. The modules that were significantly association with transpiration rate (Tr) at the *P* < 0.01 levels were marked with asterisks. B, Functional annotation of modules VuM9 and VuM17 in cowpea, and GmM4 and GmM14 in soybean.

**6 12 16	6	12 16 6 12 16	6 12 16**

**WW	MD	SD	RC**

**VuM17**

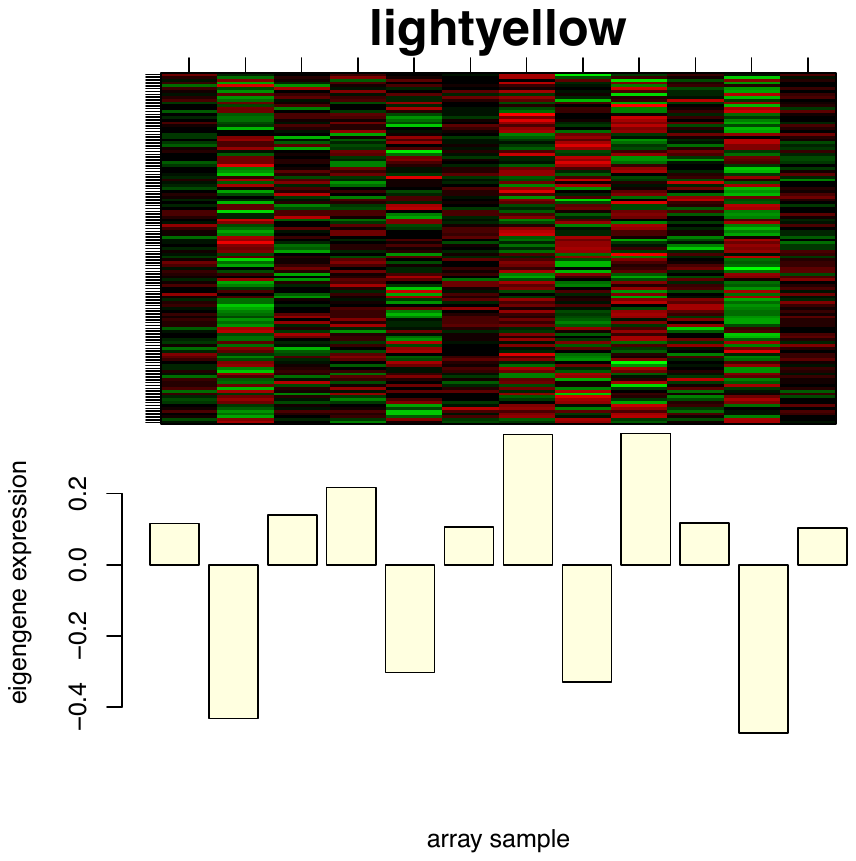

**6 12 16	6	12 16 6 12 16	6 12 16**

**WW	MD	SD	RC**

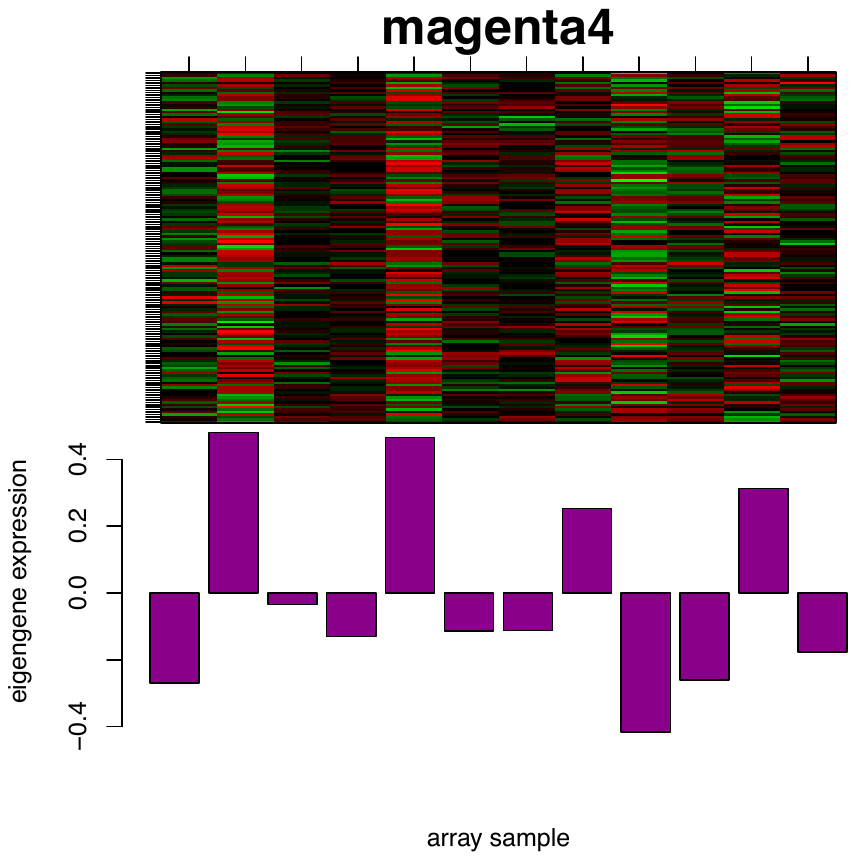

**GmM14**

**Fig. S10.** Eigengene expression of VuM17 in cowpea and GmM14 in soybean. WW: well-watered control; MD: moderate soil drought; SD: severe soil drought; RC: recovery phase. The numbers 6, 12, and 16 denote time of day (6 am, 12 pm and 4 pm).

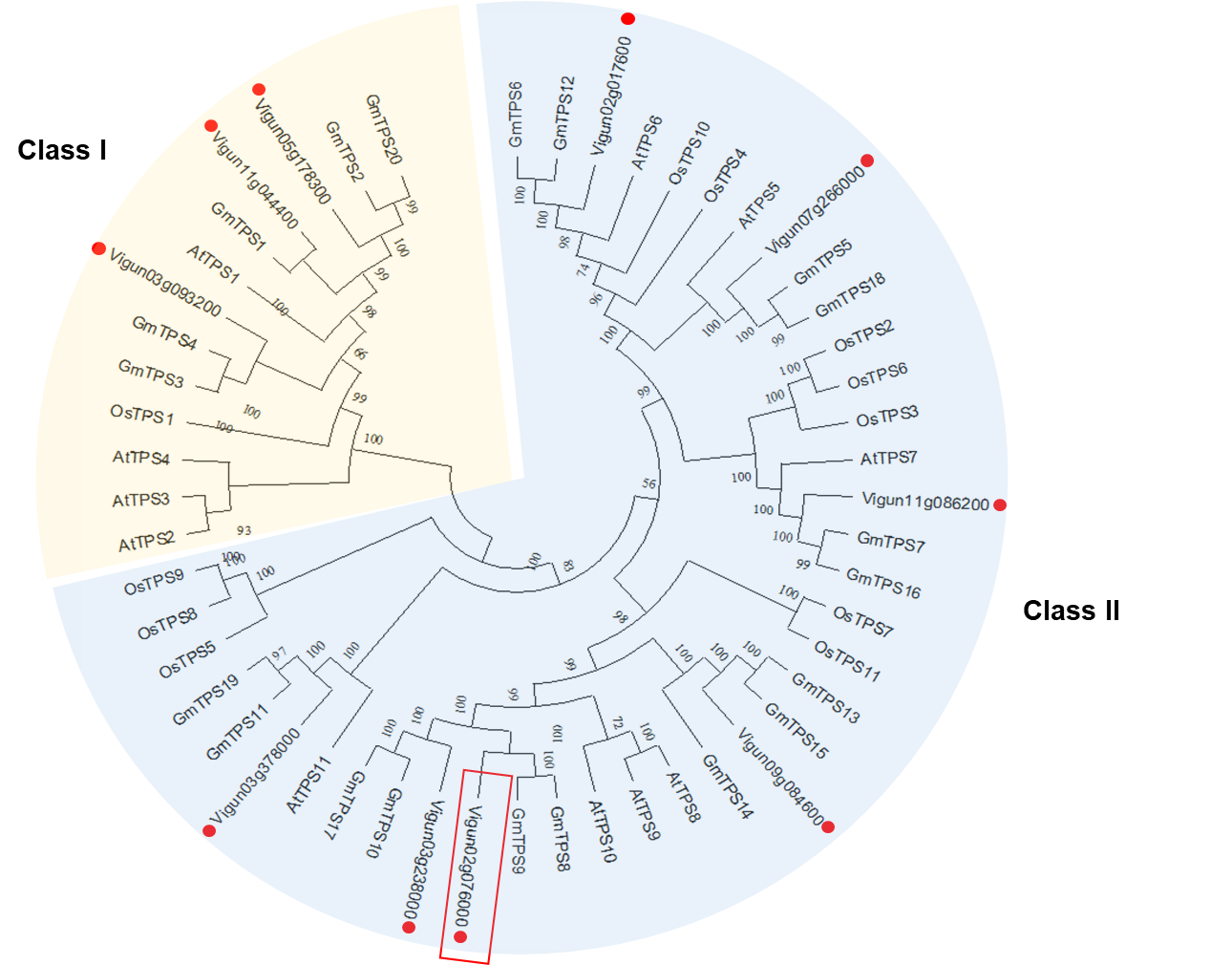

**Fig. S11.** Phylogenetic tree of the TPS proteins from *Arabidopsis thaliana*, *Oryza sativa,* *Glycine max* and *Vigna unguiculata*. The TPS proteins of *Vigna unguiculata* are marked with red circles and the *VuTPS9* is also enclosed with a red square. The phylogeny was constructed by using the maximum-likelihood method with 1000 bootstrap replicates.

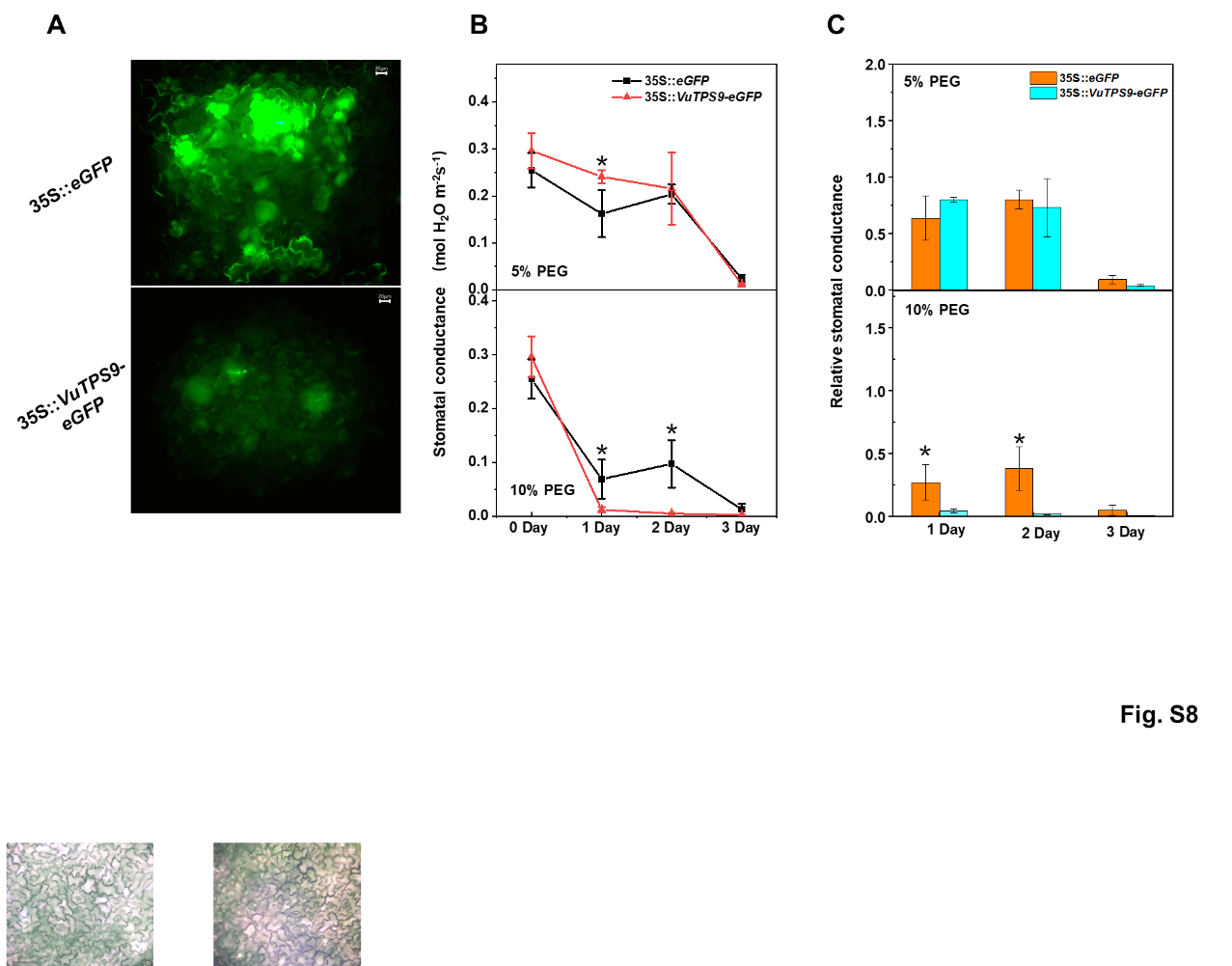
**Fig. S12.** The effect of *VuTPS9* overexpression on stomatal conductance. A, GFP expression in leaves of cowpea was detected by fluorescence microscope. Scale bars represent 20 μm. B and C, the stomatal conductance (Gs, B) and relative Gs (C) in leaves of the *VuTPS9*-OE (35S::*VuTPS9-eGFP*) and CK lines (35S::*eGFP*) after PEG-6000 treatment for 0, 1, 2 and 3 days. To calculate the relative Gs, the average Gs amount on day 0 of each line was set as 1, respectively. Data are means of at least three biological replicates with vertical error bars (±SD). Asterisk indicates significant differences according to a *t*-test at a 0.05% level of significance.
